# B cells targeting parasites capture spatially linked antigens to secure T cell help

**DOI:** 10.1101/2024.08.10.607257

**Authors:** Xin Gao, Hayley A. McNamara, Jiwon Lee, Adrian F. Lo, Deepyan Chatterjee, Dominik Spensberger, Daniel Fernandez-Ruiz, Kevin Walz, Ke Wang, Hannah G. Kelly, Kai Pohl, Patricia E. Carreira, Andrea Do, Le Xiong, Lynette Beattie, Alexandra J. Spencer, Daniel H.D. Gray, Friedrich Frischknecht, Melanie Rug, Ian A. Cockburn

## Abstract

Our understanding of T-cell-dependent humoral responses has been largely shaped by studies involving model antigens such as recombinant proteins and viruses ^1,2^. In these contexts, B cells internalize the entire antigen or pathogen, and present a range of antigens to helper CD4^+^ T cells to initiate the humoral response. However, this model does not account for large pathogens (such as parasites) that are too large to be taken up by individual B cells, and the mechanisms by which B cells acquire and present antigens from large complex pathogens to T cells remain poorly understood. Here we used *Plasmodium*, the causative parasite of malaria, as a model to investigate the requirements for T cell help for B cells targeting the *Plasmodium* surface circumsporozoite protein (CSP). Upon *Plasmodium* sporozoite (SPZ) immunization, CSP-specific B cells can form a synapse-like structure with SPZs and take up CSP and non-CSP surface antigens. As a result, CSP-specific B cells can receive help from CD4^+^ T cells specific to antigens that are located on the surface but not cytosol of the *Plasmodium* SPZ. Therefore, B cells can obtain help, not only from T cells with the same protein specificity, but also from T cells specific for spatially linked antigens. This flexibility in T cell help may enhance the initiation and maintenance of humoral immune responses to complex pathogens.

## Main

Efficient antibody responses depend upon T cell help. Helper T cells enhance the initial expansion of B cells, are critical for affinity maturation in the germinal centre (GC) and license the production of long-lived plasma cells ^3–5^. However, the majority of our understanding of T-cell-dependent humoral responses comes from studies with either model protein antigens or viruses such as influenza virus or vesicular stomatitis virus ^1,2^. In these conditions relatively few (<10) antigens are present and the probability of a B cell finding a T cell with specificity for the same protein is therefore high. However, many major pathogens are much larger, such as bacteria carrying ∼3000 genes, or eukaryotic pathogens such as the malaria parasite *Plasmodium* with ∼5000 genes ^6,7^. During immune responses to these complex pathogens it may be challenging for B cells to find T cells help targeting the same antigen specificity.

The challenge could be considerably relaxed if B cells were able to take up bystander antigens in addition to their target antigen, and thus obtain help from T cells with a range of specificities (inter- molecular help). Early study showed that priming with “shaved” influenza viruses lacking the haemagglutinin (HA) surface antigen could potentiate anti-HA immune responses to intact virions ^8^.

This provides evidence that B cells can take up small viruses and thus present T cell epitopes from several antigens to obtain inter-molecular help. However, while reptile and amphibian B cells have been reported to have significant capacity for phagocytosis there may be an upper limit to the size of antigen that can be taken up by mammalian B cells ^9,10^. In support of this study with vaccinia virus showed that preimmunization with peptides corresponding to T cell epitopes only potentiated subsequent antibody responses to the original protein and not to bystander viral proteins. Thus, it was concluded that B cells had a strong preference for help from T cells with specificity for the same protein target (intra-molecular help) ^11^.

Current malaria vaccines are based on the observation that immunization with irradiated whole *Plasmodium* sporozoites (SPZ) induces protection against live parasite challenge ^12^. Antibodies targeting the surface *P. falciparum* circumsporozoite (PfCSP) molecule are a significant component of protection leading to the development of the R21 and RTS,S subunit vaccines ^13,14^. In addition, whole parasite vaccines based on *Plasmodium* SPZ are actively being pursued. However, while SPZ-based vaccines give strong protection in malaria naïve individuals, antibody responses to PfCSP and consequently levels of protection are lower in individuals from malaria endemic areas ^15,16^. This is paradoxical, as malaria exposed individuals are expected to carry high levels of T cells specific for malaria antigens that might be expected to enhance antibody responses to SPZ ^17^.

In this study we aimed to understand how B cells acquire cognate help from CD4^+^ T cells in response to large complex pathogen. The availability of Igh^g2A10^ B cell receptor (BCR) knock-in mice in which about 2% B cells are specific for PfCSP, coupled with transgenic rodent malaria *P. berghei* parasites expressing PfCSP in place of the endogenous *P. berghei* CSP (PbCSP) provides an experimentally tractable system in which to investigate the requirements for T cell help to B cells ^18,19^. Here we show that CSP-specific B cells can obtain inter-molecular T cell help by taking up bystander antigens located on the surface of SPZs to initiate efficient antibody response. Thus, our study suggests that there is not a strict requirement for intra-molecular help during immune responses to complex pathogens.

## Results

### Pre-exposure to different Plasmodium lifecycle stages has distinct effects on the antibody response to SPZs

Our initial experiments asked whether prior exposure to malaria parasites lacking PfCSP could enhance subsequent responses to this antigen in a T-cell-dependent manner, as any enhancement observed would be evidence of inter-molecular help. Malaria parasites have a complex life cycle.

Once a SPZ enters the host, it transforms into blood stage parasites that infect red blood cells, leading to the symptomatic stage of infection. Accordingly, mice were primed with either wild type *P. berghei* SPZs (WT-SPZs) which express PbCSP and do not have PfCSP, or with WT *P. berghei* blood stage parasites (infected red blood cells, iRBC). Both sets of parasites were irradiated to arrest infection and transformation between life stages. After preimmunization mice received PfCSP-specific Igh^g2A10^ B cells and were boosted with PfCSP-expressing SPZ (PfCSP-SPZ) ^18^. Priming with iRBC was designed to mimic previous malaria exposure as seen in malaria endemic areas, while priming with WT-SPZ was designed to give the maximum opportunity for bystander enhancement of the immune response against the subsequent PfCSP-SPZ immunization, as both SPZs parasites are identical except for the CSP. Additional groups of mice received the CD4^+^ T cell depleting GK1.5 antibody prior boosting to determine if any enhancement observed was CD4^+^ T cell dependent (**Fig. 1a**).

**Figure 1.**
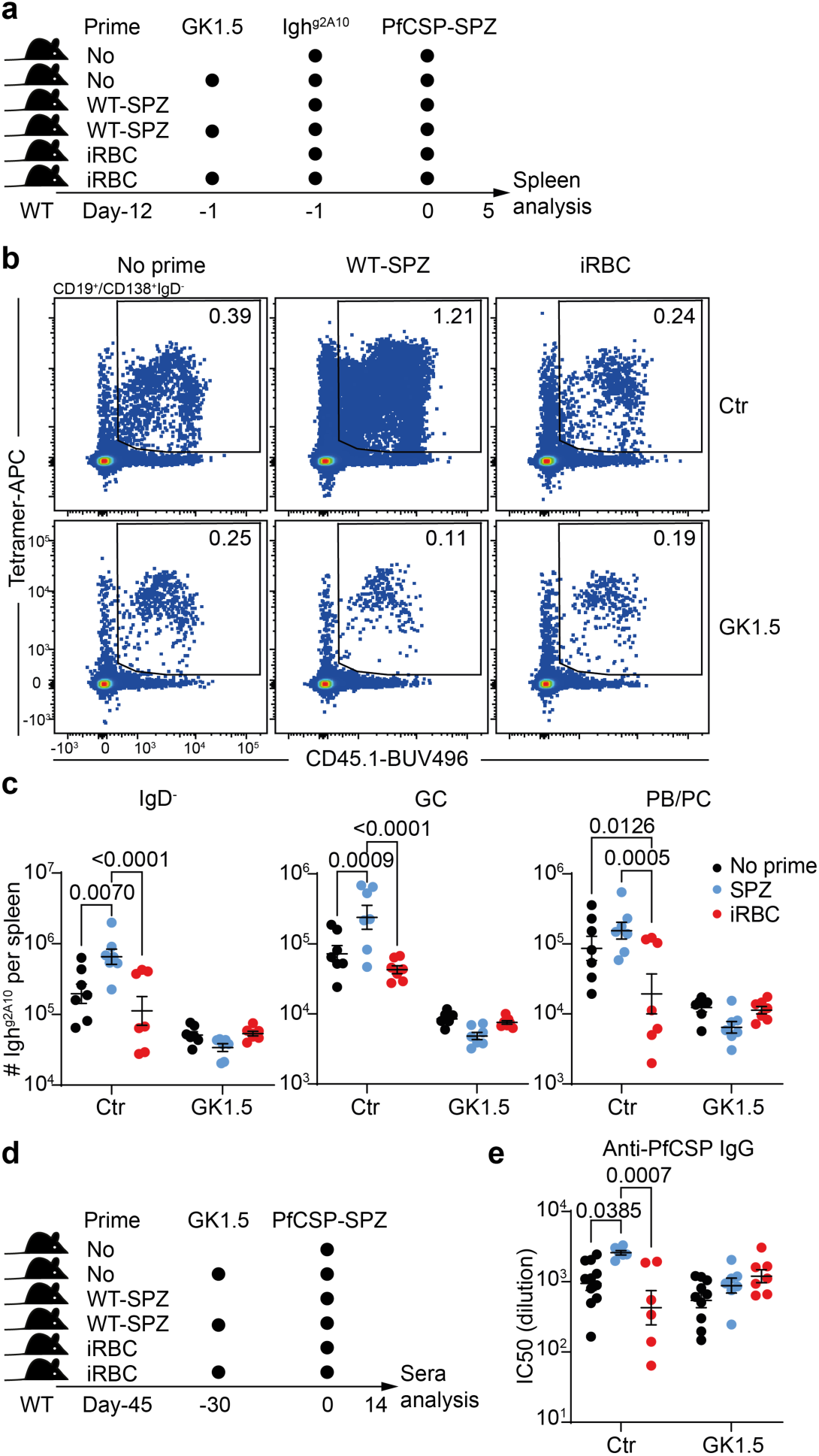
Pre-exposure to different *Plasmodium* lifecycle stages has distinct effects on the antibody response to SPZs. **a-c** , WT mice were untreated, or primed by irradiated WT-SPZ or iRBC, followed by Igh^g2A10^ B cell transfer w/wo 150μg GK1.5 treatment, and boost by PfCSP-SPZ. Spleens were analysed (n≥3). **a**, Experiment design. **b**, Representative FACS plots and **c**, statistics showing the formation of indicated Igh^g2A10^ subsets. **d-e**, WT mice were untreated, or primed by irradiated WT-SPZ or iRBC, left untreated or injected with 150μg GK1.5, followed by PfCSP-SPZ boost and sera were analysed to determine anti-PfCSP IgG (n≥3). **d**, Experiment design and **e**, statistics showing the titers of anti-PfCSP IgG of indicated groups, as measured by the IC50 of the dilution of sera. Results were pooled from two independent experiments for c, three independent experiments for e. P values were calculated by Two-Way ANOVA with Tukey multiple comparisons test for c,e.

Priming with WT-SPZ lead to significantly increased IgD^-^ (*P*=0.007, fold change (FC)=3.34) and GC (P<0.001, FC=3.31) Igh^g2A10^ B cell formation upon boosting with PfCSP-SPZ. This enhancement was CD4^+^ T cell dependent as it was not observed in GK1.5 treated animals.

Additionally, Igh^g2A10^ B cell responses were largely T-cell-dependent as GK1.5 treatment impaired these responses in all groups of mice (**Fig. 1b,c and Extended data fig. 1a**). These results indicated that WT-SPZ can act analogously to a carrier protein in a hapten carrier experiment, and boost PfCSP-specific B cell response when PfCSP is expressed on SPZ. Paradoxically, priming with iRBC did not enhance Igh^g2A10^ B cell responses following PfCSP-SPZ immunization, despite iRBC and WT-SPZ inducing comparable CD4^+^ T cell responses (**Fig. 1b,c and Extended data fig. 1a,b,c**), and theoretically expressing somewhat overlapping genes given their identical genome. To confirm if the Igh^g2A10^ B cell response correlates with antibody production, we modified the experiment by boosting the mice with PfCSP-SPZ one month after GK1.5 treatment, which allows for CD4^+^ T cell reconstitution and therefore T-cell-dependent antibody production (**Fig. 1d**). Consistent with cellular data, priming with WT-SPZ significantly enhanced (*P*=0.039, FC=2.73) while priming with iRBC had no detectable impact on anti-PfCSP IgG production (**Fig. 1e**). GK1.5 treatment eliminated the differences among all groups, confirming the enhancement was CD4^+^ T cell dependent (**Fig. 1e**).

Collectively, our results suggest that antigen-specific B cell might be able to acquire inter-molecular T cell help in the context of *Plasmodium* SPZ immunization, however it is likely that such inter- molecular help is restricted to a set of SPZ antigens that are not shared with iRBC antigens.

### Igh^g2A10^ B cells form synapse-like structures with PfCSP-SPZ

The discrepancy between the effects of pre-exposure to WT-SPZ and iRBC on subsequent responses to PfCSP SPZ suggests that antigen-specific B cell only have access to a limited range of SPZ antigens. We therefore investigated the process of B cell uptake of SPZ antigen. Accordingly, we incubated either Igh^g2A10^ or WT B cells with PfCSP-SPZs expressing the green fluorescent protein (GFP) in their cytoplasm, followed by flow cytometry, imaging flow cytometry and scanning electron microscopy (SEM) analysis (**Fig. 2a**). Parasites were stained for PfCSP, and B cells were stained for B220, and Igβ to localize the BCR. Here we found approximately 0.6% of Igh^g2A10^ B cells were associated with SPZs compared to less than 0.2% of WT cells (**Fig. 2b**), which suggests that Igh^g2A10^ B cells bind with SPZ in an antigen-specific manner. Moreover, SPZ-binding Igh^g2A10^ B cells had reduced surface PfCSP staining compared to WT B cells, reflecting the uptake of PfCSP by Igh^g2A10^ B cells (**Fig. 2c,d**). Further analysis of interactions via imaging flow cytometry revealed that while most parasites were seen resting intact on the surface of WT B cells, they were more often observed attached by only one end to Igh^g2A10^ B cells (**Fig. 2e**). Moreover, Igh^g2A10^ B cells had significantly higher co-localisation scores for PfCSP/Igβ (*P*=0.022) but not GFP/Igβ compared to WT B cells. This suggests that Igh^g2A10^ BCR can likely access PfCSP but not cytosolic GFP (**Fig. 2e,f**). Interestingly, this pattern of interaction appears to be unique to B cells: when SPZs were incubated with bone marrow derived macrophages (BMDMs) (**Extended data fig. 2a**), the SPZs were phagocytosed and their GFP was dispersed in the cytoplasm of BMDMs (**Extended data fig. 2b,c**).

**Figure 2.**
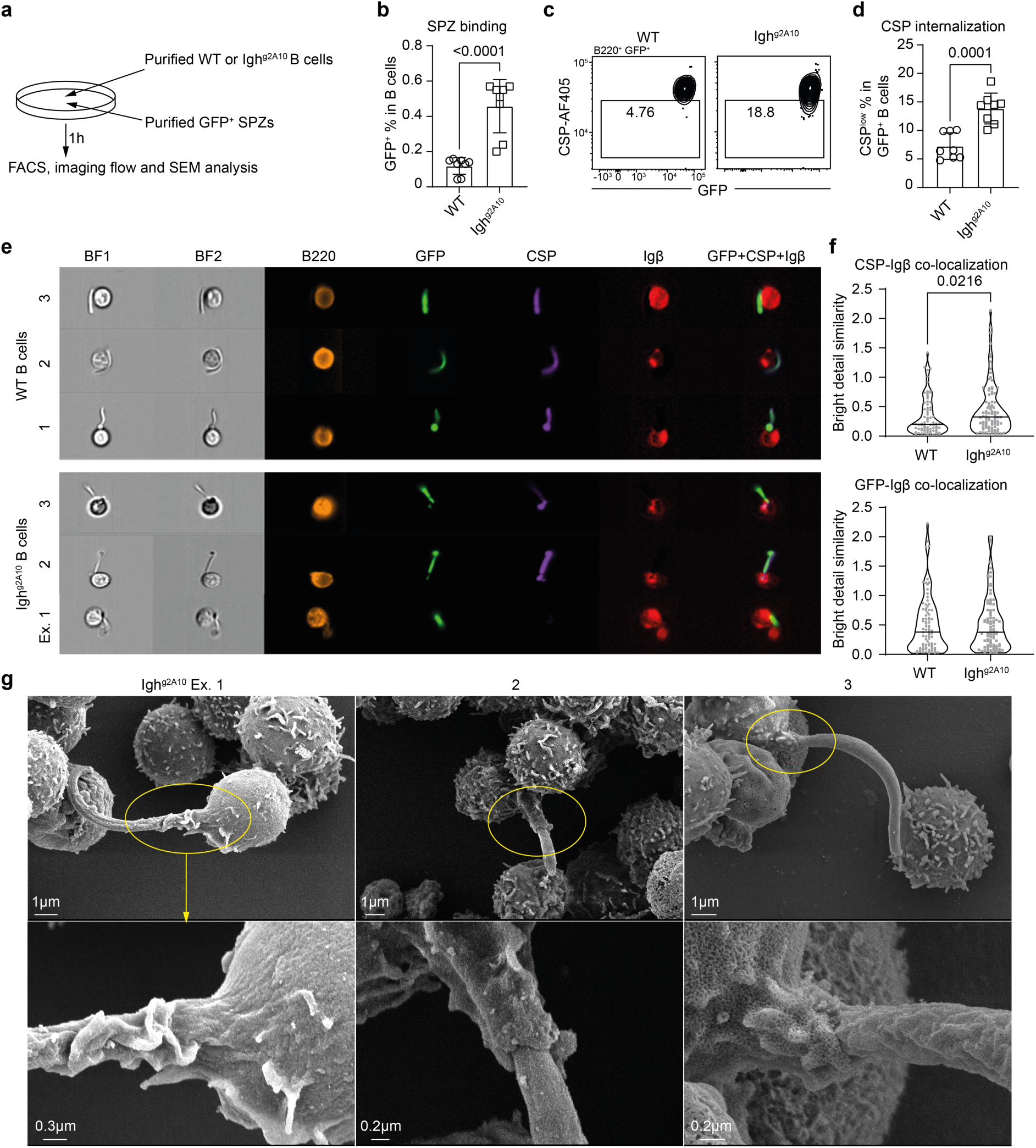
Igh^g2A10^ B cells form synapse-like structures with PfCSP-SPZ. Immunomagnetic cell separation (MACS)-enriched WT or Igh^g2A10^ B cells were incubated with FACS-purified GFP^+^ PfCSP-SPZ per well at 37℃ for 1 hour, followed by FACS, image flow or SEM analysis. **a**, Experiment design. **b**, Statistics showing the percentage of SPZ-binding B cells. **c**, Representative FACS plots and **d**, statistics showing the internalization of PfCSP by GFP^+^ B cells, indicated by the reduction of PfCSP surface staining. **e**, Representative images showing the expression of indicated markers on GFP^+^ B cells and SPZs. BF: bright field. **f**, Statistics showing the co-localization scores between indicated markers. **g**, Representative SEM images of SPZ-binding Igh^g2A10^ B cells. Lower panels show zoomed in areas indicated in yellow above. Results were pooled from three independent experiments for b, d (n≥2), two independent experiments for f, and representative of two independent experiments for g. P values were calculated by Student’s t test for b,d,f.

We further used scanning electron microscopy (SEM) to determine the nature of the interaction between SPZs and B cells. In agreement with image flow analysis, in all instances SPZ appeared only casually associated with WT B cells (**Extended data fig. 2d**). Strikingly, Igh^g2A10^ B cells appeared to have cellular membrane extrusions spread around the surface of PfCSP-SPZ, such that a proportion of the SPZ body was completely covered by B cell membrane. This pattern of tight interaction was antigen-specific, appearing in 6/15 images of Igh^g2A10^ B cells and was absent in WT B cell images. (**Fig. 2g and Extended data fig. 2e**). In summary, we showed that antigen-specific B cells can form synapse-like structures with SPZs, potentially using this structure to take up antigens.

### Direct engagement with SPZs in vivo allows optimal Igh^g2A10^ B cell responses

To determine if antigen-specific B cell-SPZ interaction also occur in vivo, we injected GFP expressing PfCSP-SPZ into either WT or Igh^g2A10^ mice to identify SPZ-binding B cells in the spleen through flow cytometry (**Fig. 3a)**. Since SPZ can bind to heparin sulphate which is abundant on many cell types ^20^, we found comparable numbers of GFP^+^ splenocytes in both WT and Igh^g2A10^ mice 1 hour post SPZ injection (**Fig. 3b,c**). We then calculated the proportions of B cells within GFP^+^ and GFP^-^ splenocytes and found that in Igh^g2A10^ mice, the B cell proportion within GFP^+^ splenocytes was significantly higher than in GFP^-^ splenocytes (*P*<0.001), whereas no such difference was observed in WT mice (**Fig. 3b,d)**. This indicates that while PfCSP-SPZs bound promiscuously to splenocytes in WT mice, they preferentially attached to B cells in Igh^g2A10^ mice. In agreement with our previous work ^21^ SPZ-associated GFP signals were undetectable after 3 hours, indicating the absence of intact SPZ from this point (**Fig. 3e)**

**Figure 3.**
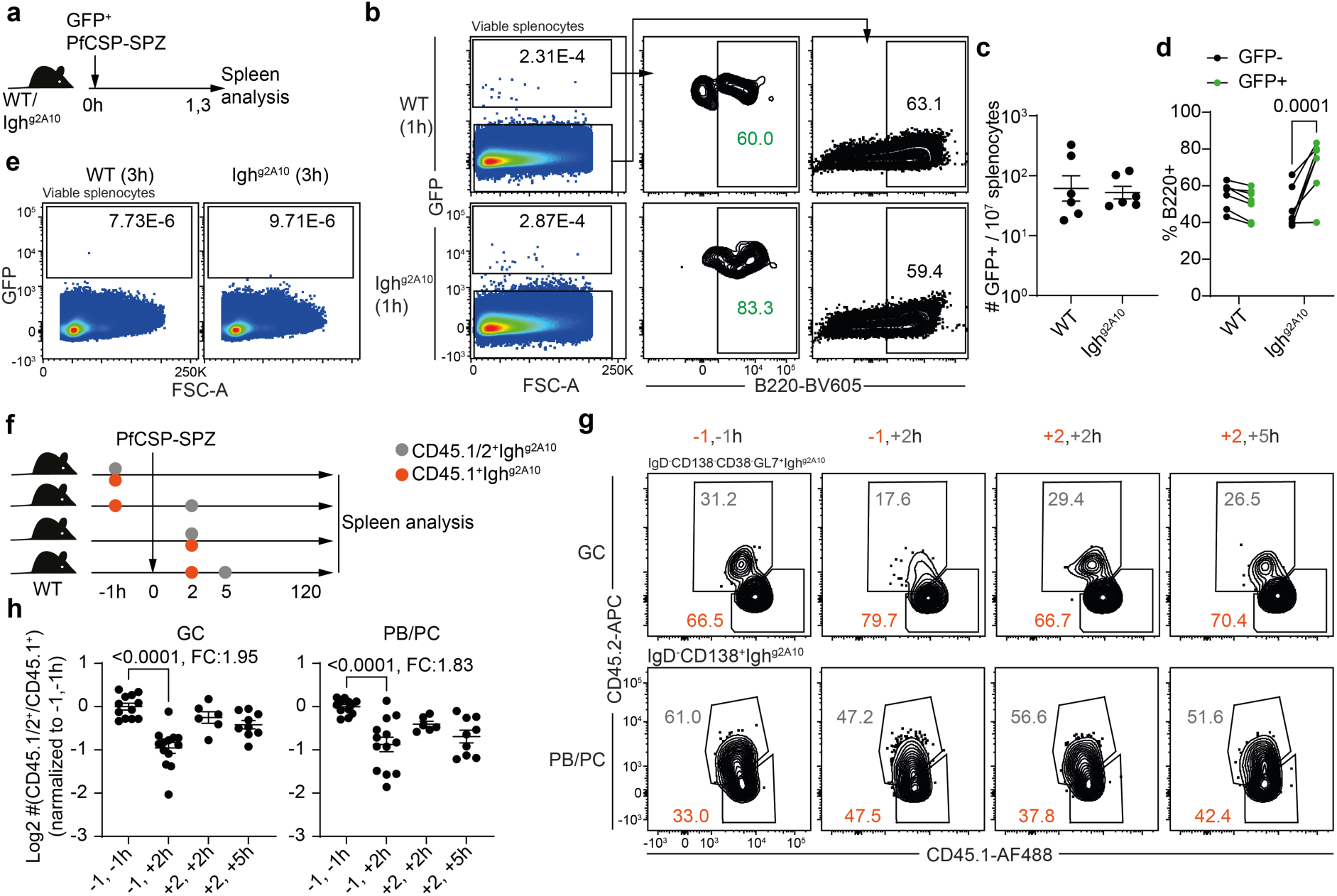
**Direct engagement with SPZs in vivo allows optimal Igh^g2A10^ B cell responses. a-e**, GFP^+^ PfCSP-SPZ were injected into WT or Igh^g2A10^ mice, and spleens were analysed after 1 or 3 hours. **a**, Experiment design. **b**, Representative FACS plots and **c**, statistics comparing the numbers of GFP^+^ cells in 10^7^ splenocytes from WT and Igh^g2A10^ mice. **d**, B220^+^ percentage in GFP^-^ or GFP^+^ splenocytes from WT or Igh^g2A10^ mice. **e**, Representative FACS plots showing the absence of GFP signal 3 hours post SPZ injection. **f-h**, PfCSP- SPZ immunized CD45.1^-^CD45.2^+^ WT mice were injected with 10^4^ CD45.1^+^CD45.2^-^ and 10^4^ CD45.1^+^CD45.2^+^ Igh^g2A10^ B cells at different timepoints, and spleens were analysed after 5 days. **f**, Experiment design. **g**, Representative FACS plots and **h**, statistics showing the formation of GC and PB/PC of CD45.1^+^CD45.2^-^ and CD45.1^+^CD45.2^+^ Igh^g2A10^ B cells transferred at indicated timepoints. Igh^g2A10^ cells were gated as CD19^+^CD45.1^+^tetramer^+^. Results were pooled from three independent experiments for c, d, h. P values were calculated by Two-Way ANOVA with Šidák multiple comparisons test for e, or with Tukey multiple comparisons test for h.

We further investigated whether such B cell-SPZ interactions contribute to B cell responses in vivo. Given that intact SPZs were absent 3 hours post injection, and B cells take minimally 1 hour to engraft into the spleen ^22^, we compared the responses of Igh^g2A10^ B cell transferred either 1 hour prior to SPZ immunization which can access intact SPZs, or 2 hours post SPZ immunization which cannot access intact SPZs (**Fig. 3f**). Under a competitive setting, Igh^g2A10^ B cells transferred 2 hours post versus 1 hour prior immunization had about a 2-fold reduction in GC and plasmablast/plasma cell (PB/PC) formation, whereas Igh^g2A10^ B cells transferred 5 hours versus 2 hours post immunization had comparable responses (**Fig. 3g,h**). This suggests that failing to engage with the intact SPZs, rather than the delayed transfer, significantly impaired PfCSP-specific B cell responses to SPZ. Collectively these data indicate that antigen-specific B cells could directly bind with whole SPZs in vivo, and such early engagement was required for the B cells to display optimal responses in a later stage.

### Limited TCR diversity in Igh^g2A10^ -helping Tfh cells during SPZ immunization

In our previous experiment we found that Igh^g2A10^ B cells could attach to the surface of SPZs and thus potentially take up surface antigens, however it remained unclear whether Igh^g2A10^ B only acquired PfCSP, or other bystander antigens during this process. To investigate this we aimed to study the T cell receptor (TCR) repertoire of T follicular helper (Tfh) cells that provide help to Igh^g2A10^ B cells upon SPZ immunization. We designed an experiment in which Igh^g2A10^ cells were transferred to MD4 mice that only carry B cells specific for hen egg lysozyme (HEL) and immunized with either recombinant PfCSP in alum (rPfCSP-alum, CSP group) or PfCSP-SPZ (SPZ group). Subsequently Tfh cells (CXCR5^+^PD-1^+^) were sorted for single-cell RNA-seq (scRNA-seq)/TCR-seq. MD4 mice were chosen as recipients to ensure all Tfh help was directed towards the transferred Igh^g2A10^ B cells. We reasoned that any bystander antigen uptake in the SPZ group would result in a more diverse TCR repertoire compared to the CSP group. As controls we also included Tfh cells from unimmunized MD4 mice (UT group), and PfCSP-SPZ immunized MD4 mice but with no Igh^g2A10^ B cell transfer (NoB group) (**Fig. 4a and Extended data fig. 3a,b**).

**Figure 4.**
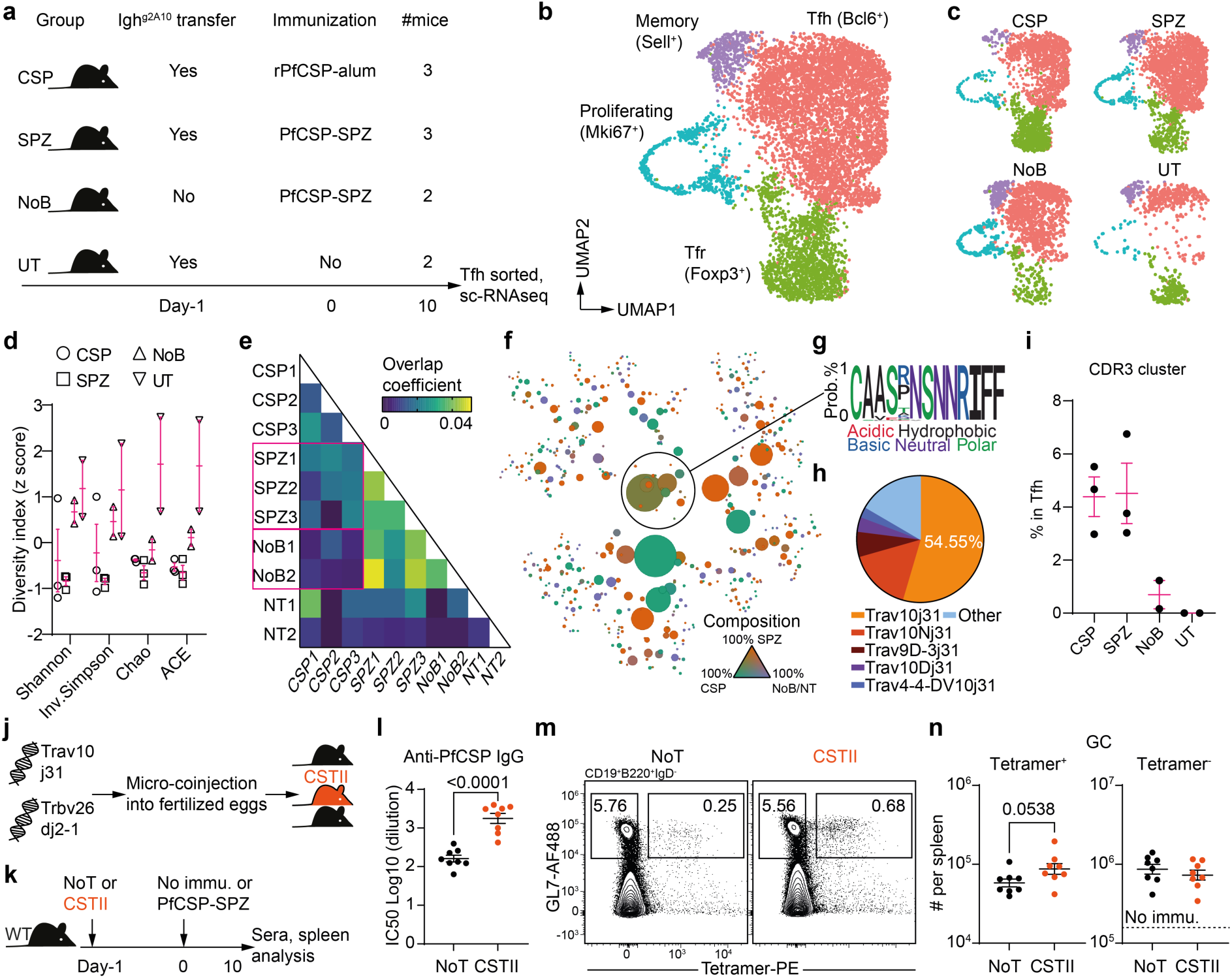
**Limited TCR diversity in Igh^g2A10^ -helping Tfh cells during SPZ immunization a-i**, MD4 mice were untreated or transferred with Igh^g2A10^ B cells, followed by rPfCSP-alum, PfCSP-SPZ immunization or left untreated, and spleens were collected. Tfh cells (CD4^+^CD44^+^PD-1^+^CXCR5^+^) were FACS- purified, hashtaged and pooled from scRNA-seq. **a**, Experiment design. UMAPs of **b**, all CD4^+^ T cells and **c**, CD4^+^ T cells separated by treatment groups. **d**, Clonal diversity for each group calculated by indicated algorithms. **e**, Heatmap of TCR clonal overlap coefficient among groups. **f**, Bubble plot showing all TCR clones based on the CDR3 amino acid sequences of TCRα. Each circle indicates a unique CDR3 sequence, and the size of the circle indicates clonal size. The proximity of different circles indicates the similarity of CDR3 amino acid sequences. The color indicates the clonal composition. **g**, Consensus amino acid sequence of circled CDR3 cluster. **h**, TCR α chain V and J gene usage of the CDR3 cluster. **i**, Relative clonal abundance of the CDR3 cluster in Bcl6^+^ Tfh cells in each mouse. **j**, Experiment design of generating TCR transgenic CST-II mice derived from the most expanded TCR sequences of highlighted CDR3 cluster. **k-n**, WT mice were untreated or transferred with 10^5^ CST-II CD4^+^ T cells, and immunized with PfCSP-SPZ. Sera and spleens were analyzed. **k**, Experiment design. **l**, Statistics showing the titers of anti-PfCSP IgG. **m**, Representative FACS plots and **n**, statistics showing the numbers of PfCSP-specific and non-specific GCB cells. Results were pooled from two independent experiments for l,n. P values were calculated by Student’s t test for l,n.

Uniform manifold projection (UMAP) analysis of transcriptomic data revealed expected clusters of Bcl6^+^ Tfh and Foxp3^+^ T follicular regulatory (Tfr) cells in all mice, as well as a small number of CD62L^+^ memory cells and a population of Ki67^+^ proliferating cells (**Fig. 4b and Extended data fig. 3c**). Re-clustering on Bcl6^+^ Tfh cells resulted in Tfh precursors (PreTfh) and several GC Tfh subsets expressing signature genes including Hif1α, IL-21 and Ascl2 (**Extended data fig. 3d,e**) ^23,24^. As expected, CSP and SPZ groups had more Tfh especially GC Tfh cell expansion than control groups (**Fig. 4c and Extended data fig. 3f**).

We then calculated the TCR diversity index in each group to look for evidence for bystander antigen uptake which should result in inter-molecular help. As predicted, the UT group had the highest TCR diversity. To our surprise, CSP and SPZ groups had comparable diversity, suggesting that Igh^g2A10^ B cells were predominately helped by PfCSP-specific Tfh cells in the SPZ group (intra- molecular help) (**Fig. 4d**). In line with this, overlapping clones were more common in CSP/SPZ groups compared to CSP/NoB groups (**Fig. 4e**). Further immune receptor repertoires clustering of expanded clones revealed a GC Tfh-featured clone family in CSP and SPZ groups, with consensus CDR3 sequence CAASXNSNNRIFF and Trav10-Trbv16/26 as the dominant variant genes (**Fig. 4f,g,h and Extended data fig. 3g,4a**). The proportion of this clone family in Tfh cells were comparable in CSP and SPZ groups, further indicating that PfCSP-specific B cells preferentially obtain intra-molecular help in the context of whole parasite immunization (**Fig. 4i**).

To formally determine whether the clone family we identified was in fact PfCSP-specific, we generated TCR transgenic mice designated CST-II that express the dominant TCR α and β (Trav10j31-Trbv26j2-1) of this clone family (**Fig. 4j and Extended data fig.4b,c**). As predicted, CST-II CD4^+^ T cells specifically expanded in response to rPfCSP-alum and to PfCSP-SPZ, but not to WT-SPZ immunizations (**Extended data fig. 4d,e,f**). Epitope mapping further identified a 16mer from the major repeat (NANP)4 as the cognate epitope for the CD4^+^ T cells (**Extended data fig. 4g,h,i,j,k**). Lastly, we showed that CST-II cells can efficiently help Igh^g2A10^ B cells responses in PfCSP-SPZ immunization (**Extended data fig. 4l,m)**.

To confirm if B cells preferentially utilise intra-molecular T cell help in SPZ immunization, we asked whether adoptive transfer of CST-II cells could potentiate overall B cell responses to PfCSP-SPZ, or only responses to PfCSP (**Fig. 4k**). We found that antibody responses to PfCSP were significantly enhanced, as was the number of PfCSP-specific GC B cells, however the overall magnitude of the GC response was not altered when we augmented the CSP specific T cell response via the addition of CST-II cells (**Fig. 4l,m,n**). In summary, these results suggest that PfCSP-specific B cells are preferentially helped by PfCSP-specific CD4^+^ T cells in primary PfCSP-SPZ immunization. These results are in agreement with previous experimental data using vaccinia virus which suggests that CD4^+^ T cell help to B cells was nontransferable to other virion protein specificities ^11^.

### PfCSP-specific B cells can obtain inter-molecular help during SPZ immunization

Intriguingly, our WT-SPZ priming data indicates PfCSP-specific B cells can access multiple SPZ antigens and obtain inter-molecular help. However, the TCR-seq data suggests that PfCSP-specific B cells preferentially take up PfCSP and acquire intra-molecular help. To reconcile the conflicting results, we asked if PfCSP-specific B cell can still respond to SPZ immunization in the absence of PfCSP-specific T cell help (intra-molecular help). We developed mice which carried functional PfCSP-specific B cells but whose T cells are tolerant to PfCSP. This was achieved by generating mCherry-rev-mPfCSP-fl mice which express double-floxed mCherry allele under the control of a CAG promoter upstream of a PfCSP (transmembrane domain-fused, mPfCSP) in the reverse orientation. In these mice when Cre is expressed, the mCherry allele is excised and replaced by the mPfCSP gene in the correct orientation (**Fig. 5a**). Crossing of these mice to a Foxn1 Cre line (PfCSP^T-tol^) resulted in the expression of mPfCSP in thymic epithelial cells (TEC) of the progeny mice, and thus the tolerance of T cells but not B cells towards PfCSP. Additionally, mice were crossed to a ubiquitous germline PGK Cre (PfCSP^full-tol^) to allow for germline transmission and expression of PfCSP in all tissues, resulting in mice developed full tolerance (both T and B cells) to PfCSP (**Fig. 5b**). We confirmed the loss of mCherry and the expression of PfCSP only in the TEC of PfCSP^T-tol^ mice, and in all cells in PfCSP^full-tol^ mice (**Fig. 5c,d**).

**Figure 5.**
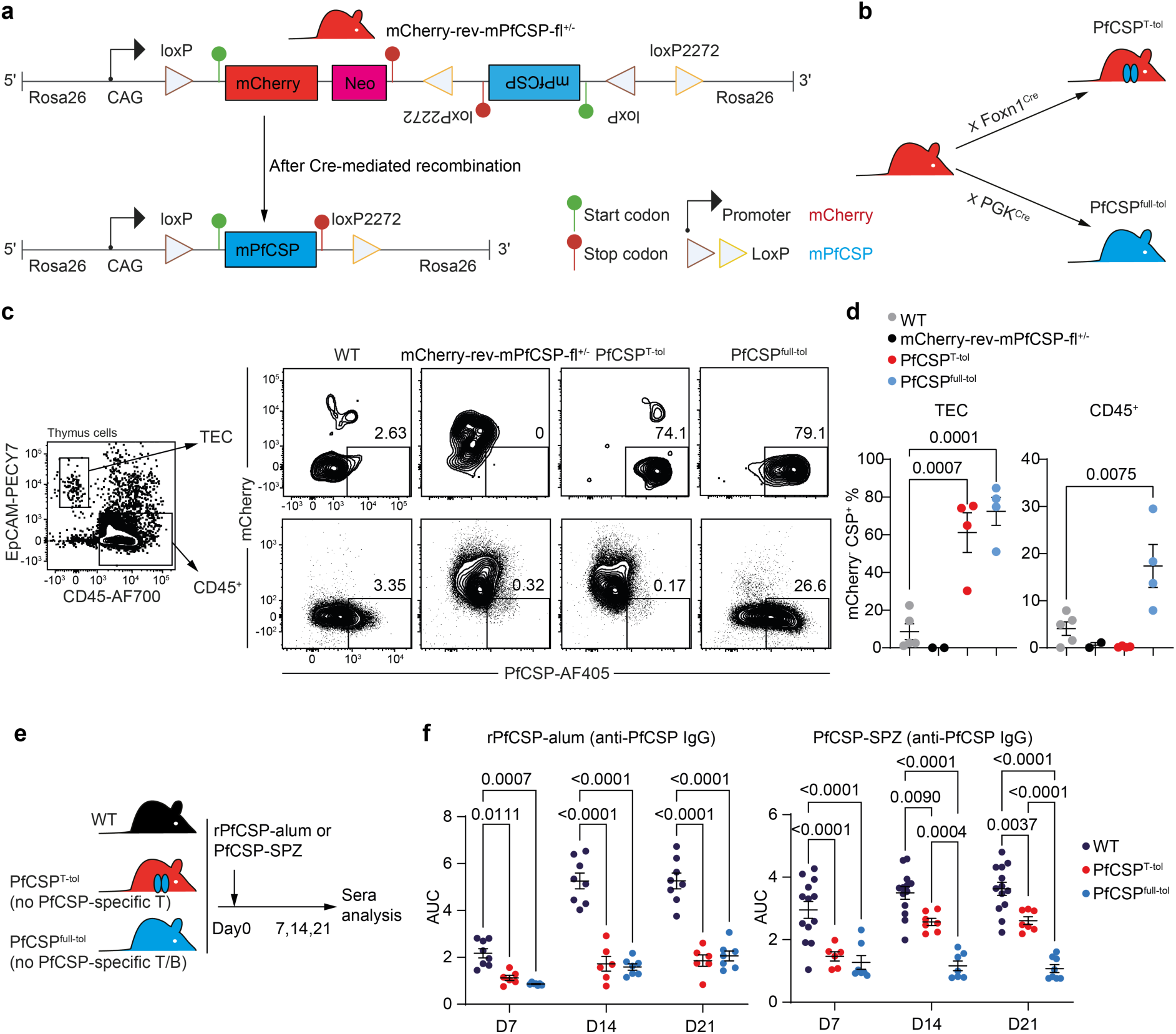
**PfCSP-specific B cells can obtain inter-molecular T cell help during SPZ immunization. a**, Diagram showing the DNA constructs for generating mCherry-rev-mPfCSP-fl knock-in mice, and the result after Cre-mediated recombination. **b**, Breeding strategy to create PfCSP^T-tol^ (mCherry-rev-mPfCSP-fl^+/-^ Foxn1cre^+/-^) and PfCSP^full-tol^ (mCherry-rev-mPfCSP-fl^+/-^ PGKcre^+/-^) mice. **c**, Representative FACS plots and **d**, statistics showing the expression of mCherry and intracellular PfCSP on indicated thymus cell subsets (n≥1). **e-f**, WT, PfCSP^T-tol^, and PfCSP^full-tol^ mice were immunized by rPfCSP-alum or PfCSP-SPZ, and anti-PfCSP IgG were analysed. **e**, Experiment design and **f**, statistics showing the titers of anti-PfCSP IgG under indicated conditions (n≥3). Results were pooled from ≥two independent experiments for d,f. P values were calculated by One-Way ANOVA with Dunnett multiple comparisons test for d, and with Tukey multiple comparisons test for f.

To determine whether functional PfCSP-specific CD4^+^ T cells were deleted in either PfCSP^T-tol^ or PfCSP^full-tol^ mice, we generated mixed bone marrow chimeric mice by transferring 50% CST-II/50% wild type (WT) bone marrow cells into either PfCSP^T-tol^ or PfCSP^full-tol^ animals (**Extended data fig. 5a**). Unexpectedly in these conditions, CST-II cells were only partially deleted in the thymus in both groups, and even proliferated in the spleen of the PfCSP^full-tol^ mice potentially due to their activation in the periphery (**Extended data fig. 5b,c,d,e**). Similar results were obtained by crossing these lines to CST-II background (**Extended data fig. 5f,g)**. Nevertheless, the residual CST-II CD4^+^ T cells in these crossed animals appeared to be anergic because they were not able to respond to rPfCSP-alum immunization after adoptive transfer into WT animals (**Extended data fig. 5h,i,j)**.

Similarly, PfCSP^T-tol^ and PfCSP^full-tol^ mice were crossed to Igh^g2A10^ mice to determine whether they can support the development of functional PfCSP-specific B cells. PfCSP-specific B cells were expanded in PfCSP^full-tol^ mice but possessed features of anergy ^25^ including elevated IgD and reduced IgM, Igβ and Igκ, while these features were not observed in PfCSP^T-tol^ mice (**Extended data fig. 6a,b,c,d**). In agreement with this adoptively transferred B cells from crossed PfCSP^full-tol^ mice were unable to respond to rPfCSP-alum immunization whereas B cells from crossed PfCSP^T-tol^ mice were able to respond (**Extended data fig. 6e,f,g**). Collectively, we confirmed that PfCSP^T-tol^ mice only lacked CD4^+^ T cells capable of responding to PfCSP, while PfCSP^full-tol^ mice lacked both functional CD4^+^ T cells and B cells specific to PfCSP.

**Figure 6.**
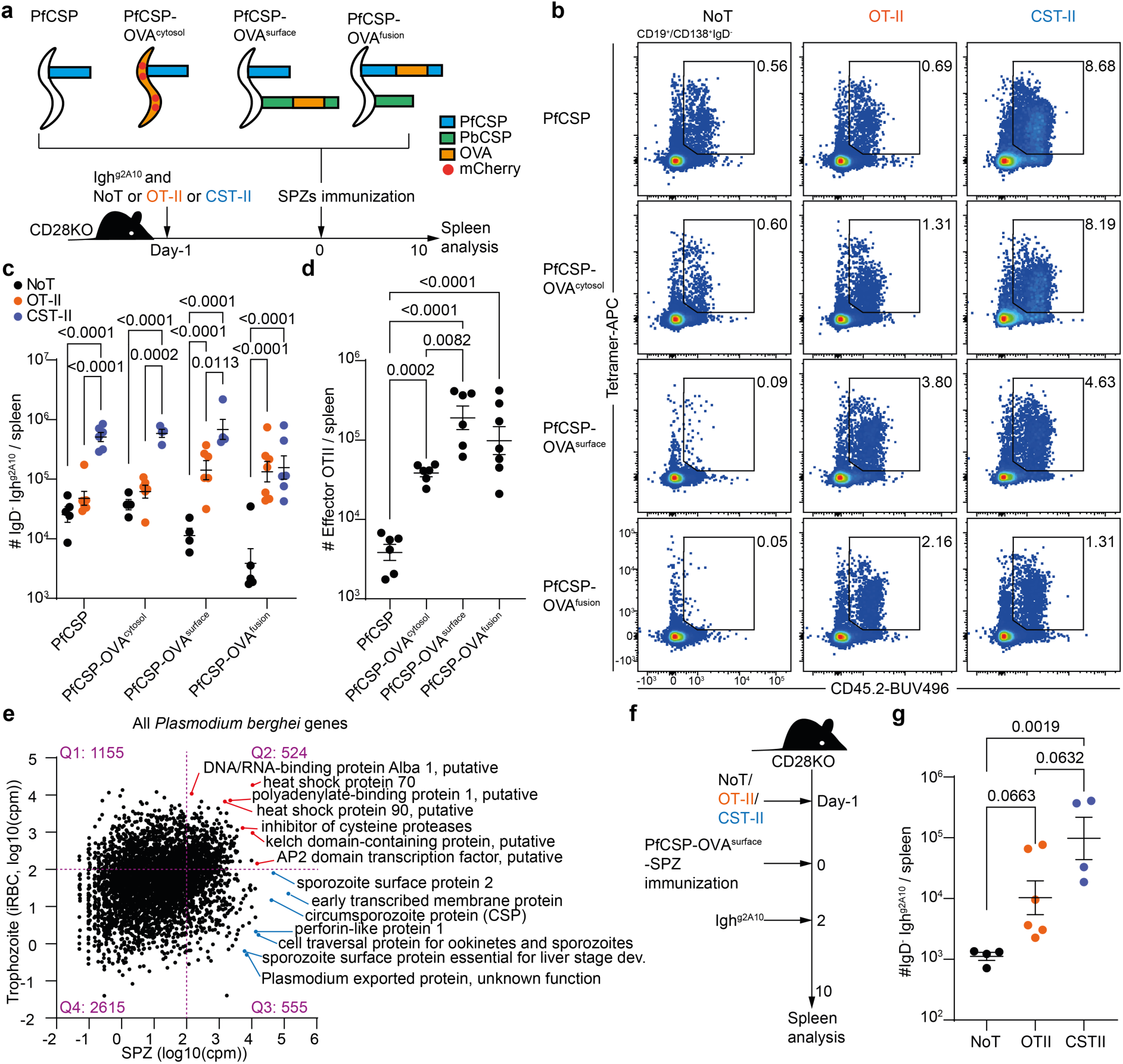
Inter-molecular help for PfCSP-specific B cells is provided by CD4^+^ T cells specific to SPZ bystander surface antigens. a-d. , CD28^-/-^ mice were transferred with Igh^g2A10^ B cells w/wo OT-II or CST-II CD4^+^ T cells, followed by immunization with transgenic SPZs, and spleens were analysed. **a**, Experiment design (n≥2). **b**, Representative FACS plots and. **c**, statistics showing the formation of IgD^-^ Igh^g2A10^ B cells under indicated conditions. **d**, Statistics showing the numbers of effector OT-II CD4^+^ T cells after indicated immunizations. **e**, mRNA expression of *P. berghei* genes in the trophozoite (GSE252704) and SPZ (GSE212753) stages. Count per million (cpm)=100 is the threshold to distinguish lowly and highly expressed genes. **f-g**, CD28^-/-^ mice were transferred w/wo OT-II or CST-II CD4^+^ T cells, immunized by PfCSP-OVA^surface^-SPZ, transferred with Igh^g2A10^ B cells 2 days after immunization, and spleens were analysed (n≥2). **f**, Experiment design. **g**, Statistics showing the formation of IgD^-^ Igh^g2A10^ B cells. Results were pooled from three independent experiments for b,d, and two independent experiments for g. P values were calculated by Two-Way ANOVA with Tukey multiple comparisons test for c, and by One-Way ANOVA with Tukey multiple comparisons test for d,g.

Having established that PfCSP^T-tol^ and PfCSP^full-tol^ mice behaved as expected we immunized these mice and WT littermates with either rPfCSP-alum or PfCSP-SPZ, to formally test whether CSP-specific B cells only receive intra-molecular help, or also receive inter-molecular help during SPZ immunization (**Fig. 5e**). As expected upon rPfCSP-alum immunization, only WT mice were able to mount anti-PfCSP IgG, as in this setting efficient antibody response requires both PfCSP-specific B and T cells. However, when PfCSP-SPZ were used for immunization, PfCSP^T-tol^ but not PfCSP^full-tol^ mice were also able to mount anti-PfCSP IgG responses (**Fig. 5f**), suggesting that in this context PfCSP-specific B cells were able to access multiple SPZ antigens and obtain inter-molecular help from non-PfCSP specific CD4^+^ T cells.

### Inter-molecular help for PfCSP-specific B cells is provided by CD4^+^ T cells specific to SPZ bystander surface antigens

Our final question was to identify the origin of inter-molecular help. Our SEM imaging data suggests that the inter-molecular help came from the surface but not cytosolic SPZ antigens (**Fig. 2g**). First, to test if CSP-specific B cells can acquire inter-molecular help from T cell specific for cytosolic antigens, we transferred Igh^g2A10^ B cells with irrelevant OT-II cells, or PbT-II cells specific for *P. berghei* cytosolic heat shock protein 90 (Hsp90) ^26^ into CD28-/- mice followed by PfCSP-SPZ immunization (**Extended data fig. 7a**). In this setup, help only comes from the transferred TCR transgenic cells as CD28-/- mice do not develop functional Tfh cells ^27^. We found PbT-II cells significantly expanded under immunization, but the numbers of IgD^-^ Igh^g2A10^ B cells were comparable in OT-II and PbT-II transferred mice (**Extended data fig. 7b,c**). This suggests that CSP-specific B cells do not acquire inter-molecular help from cytosolic antigens. To further determine if the inter- molecular help comes from bystander surface antigens, we transferred CST-II or irrelevant OT-II cells into CD28-/- mice and immunized them with transgenic SPZs expressing PbCSP and PfCSP as separate proteins on the SPZ surface ^28^ (**Extended data fig. 7d**). In this context we found CST-II cells, which are specific to PfCSP, significantly enhanced antibody response to PbCSP (**Extended data fig. 7e,f**). This suggests that CSP-specific B cells can acquire inter-molecular help from bystander surface antigens.

In previous experiments, different transgenic T cells might have intrinsically different capacity to help B cells (e.g. different TCR affinity). To address this limitation, we generated three lines of transgenic parasites co-expressing PfCSP and the OVA323-339 (OT-II epitope) in different relative spatial locations. First, PfCSP-OVA^cytosol^ contained PfCSP on the surface of the SPZ and OVA (whole protein, fused with mCherry) expressed cytosolically under the control of a strong constitutive promoter Hsp70 (**Extended data fig. 8a**). Second, PfCSP-OVA^surface^ carried intact PfCSP, and PbCSP molecule with OVA260-386 inserted between the repeat and C terminus in order to target OVA to the sporozoite surface but on a separate protein (**Extended data fig. 8b**). Finally, PfCSP-OVA^fusion^ carried OVA260-386 inserted between the repeat and C-terminus of PfCSP. Since the insertion of the OVA260-386 rendered PfCSP non-functional it was also necessary to insert PbCSP to ensure SPZs can still emerge from mosquitoes (**Extended data fig. 8c**). The expression of transgenes on SPZs was validated via flow cytometry (**Extended data fig. 8d**).

We used these parasites to immunize CD28^-/-^ mice that had received Igh^g2A10^ cells and either no T cells (negative control), CST-II cells (positive control) or OT-II cells (experimental group) to determine the source of inter-molecular help to Igh^g2A10^ B cells (**Fig. 6a**). Our readout in this experiment was the number of Igh^g2A10^ cells responding to PfCSP. As expected, we detected only a small number of IgD^-^ Igh^g2A10^ cells in CD28^-/-^ mice without transferred T cells while CST-II cells were able to support Igh^g2A10^ B cell responses in immunizations with all strains of SPZ (**Fig. 6b,c**).

Interestingly, OT-II cells were unable to help when OVA was not present, or was present in the SPZ cytosol. However, OT-II cells significantly supported Igh^g2A10^ B cells responses when OVA was fused within PfCSP (PfCSP-OVA^fusion^) (P<0.001, FC=34.4), or present on the SPZ surface even when not inserted into PfCSP itself (PfCSP-OVA^surface^) (P<0.001, FC=12.6) (**Fig. 6b,c**). Similar results were observed for Igh^g2A10^ GC B cell and PB/PC formation (**Extended data fig. 9a**). Importantly OT-II cells significantly expanded in response to all strains of OVA-expressing SPZ (P<0.001), although the magnitude was lower for PfCSP-OVA^cytosol^-SPZ (**Fig. 6d and Extended data fig. 9b**). In sum these results showed that Igh^g2A10^ B cells can receive help from CD4^+^ T cells that are specific for bystander surface, but not cytosolic SPZ antigens.

Our previous experiment demonstrated that inter-molecular help comes from bystander surface antigens, however it was confounded by the different magnitudes of OT-II expansion. We therefore performed an analogous experiment in which OT-II cells were transferred to MD4 mice with or without Igh^g2A10^ cells present (**Extended data fig. 9c**). Because Tfh formation requires antigen presentation from B cells ^29^, in this set up we would expect OT-II Tfh cell formation if the Igh^g2A10^ cells are able to take up OVA from the different strains of SPZ. Analogously to results in CD28^-/-^ mice, OT-II Tfh cells were only formed in MD4 mice transferred with Igh^g2A10^ cells and immunized with SPZ expressing OVA on the surface, but not in cytosol (**Extended data fig. 9d,e**). Of note, the lack of Tfh formation in PfCSP-OVA^cytosol^-SPZ immunized mice was not due to poorer Igh^g2A10^ GC B cell formation (**Extended data fig. 9f,g**), nor less OVA availability, as we observed comparable OT-II effector cell formation between OVA^cytosol^ and ^surface^ parasites without Igh^g2A10^ cell transfer (**Extended data fig. 9h,i**).

Our findings may also provide an explanation for the paradox that WT-SPZ, but not iRBC priming can enhance Igh^g2A10^ responses to PfCSP-SPZ (**Fig. 1a-e**). Analysis of the transcriptomes of trophozoites (iRBC) and SPZ revealed that shared transcripts are enriched only for cytosolic proteins such as chaperones and transcription factors ^30^. In contrast the genes that are unique to SPZ are highly enriched for surface proteins (**Fig. 6e**). We also took another approach by analysing antigens identified on the surface of *P. falciparum* SPZ by mass spectroscopy ^31^. Among the 32 homologous genes putatively located on the surface of *P. berghei* SPZ, we found their overall expression in SPZs were negatively correlated with their expression in iRBC (**Extended data fig. 9j**). Such gene expression pattern further indicated that iRBCs lack the antigens that are abundant on the surface of SPZ, thus prior blood stage infection cannot provide inter-molecular help to enhance CSP-specific B cell responses.

Finally, because B cell can not only acquire antigens directly from the SPZ, but also from other sources like subcapsular sinus macrophages (SCSMΦ) and follicular dendritic cells (FDCs) ^32,33^, we asked whether inter-molecular help was possible even once intact SPZ were no longer present in the spleen. Accordingly, we immunized CD28^-/-^ mice that had received no T cell transfer, OT-II or CST-II cells with PfCSP-OVA^surface^-SPZ, but waited two days to ensure the absence of intact SPZs before transferring Igh^g2A10^ cells into these animals (**Fig. 6f**). We found even in these circumstances OT-II cells were able to provide inter-molecular help to Igh^g2A10^ B cells, although their help was inferior to that from CST-II cells (**Fig. 6g and Extended data fig. 9k**). These results indicated that SPZ antigens are likely maintained by the immune system as complexes comprised of multiple protein molecules which are available for uptake by B cells.

## Discussion

In this study we aimed to investigate the antigenic requirements for T cell help in antibody response towards large complex pathogens. We showed that antigen-specific B cells can directly engage with SPZs and take up cognate and bystander surface antigens for presentation, obtaining both intra-molecular and inter-molecular T cell help. Our finding partially contradicts a previous study showing that priming with predicted T cell epitopes from vaccinia virus antigens enhanced antibody responses to that antigen alone, rather than to other virus antigens ^11^. Similarly, our scRNA-seq analysis also found Igh^g2A10^ B cells mainly obtained intra-molecular T cell help in the SPZ immunization. Providing that Tfh cells are selected by B cells in a GC reaction ^34^, and intra-molecular help is generally more efficient that inter-molecular help (as shown in our PfCSP^T-tol^ mice), one interpretation for the discrepancy is that after multiple rounds of selection in the GC, Tfh cells have “evolved” to share the same specificity with B cells. Therefore, it is possible that inter-molecular help would be more pronounced in the early stages of B cell responses, perhaps ensuring that B cells can find T cell help in a highly diverse TCR environment, while intra-molecular help dominates the later responses to ensure the stringent selection in the GC. As such, it would be interesting to compare the helper function of OT-II versus CST-II cells to PfCSP-specific B cells at different timepoints in response to SPZ immunization. Additionally, GC B cells use distinct molecular pathways to extract antigens compared to naïve and memory B cells ^35^, so it is possible that the ability to acquire inter- molecular help for a GC B cell is different from a naïve B cell.

We further show that CSP-specific B cells can still acquire inter-molecular help in the absence of intact SPZs. This implies that SPZ antigens are retained as multi-protein complexes by the immune system over an extended period of time. It is currently not clear whether parasitic proteins that are in close proximity to each other more likely to be stored in the same complex by the immune system, or whether all parasitic antigens are presented randomly (e.g. on the surface of FDCs). Previous studies showed that SCSMΦ can process foreign antigens, and antigen-specific B cells transfer these antigens (as immune complexes) from SCSMΦ to FDCs ^36^, which then retain the antigens for a long period of time through endocytosis and recycling ^37^. It would be interesting to test if this antigen presentation pathway also applies to SPZ immunization, and if so, how different SPZ antigens are stored in SCSMΦ and FDCs.

We showed that antigen-specific B cells can form the synapse-like structure with SPZ. This pattern mimics the way B cells acquire antigens from antigen-loaded cells through immune synapse formation ^38^. A follow up question would be to untangle the molecular mechanism that involves in such B cell-SPZ interaction. Previous studies have showed that B cells secrete lysosomes to facilitate antigen uptake from anti-BCR coated beads ^39^, and B cells have been showed to use mechanical force to pull antigens from the plasma membrane sheets ^40^. Moreover, clathrin, endophilin and caveolin have been showed to participate in B cell antigen uptake ^41^. It would be interesting to test if these mechanisms also apply to antigen uptake from large complex pathogens. Additionally, in GC higher affinity B cells are typically more competitive for T cell help ^42^. However, it remains unclear if higher BCR affinity can also result in capturing more bystander antigens and gaining more inter-molecular help. It is also possible that very high affinity BCR may only capture the cognate antigen, resulting in less membrane spreading, fewer bystander antigen capturing and poorer ability to obtain inter- molecular help. Therefore, it would be interesting to generate BCR knock-in mice with different affinities towards PfCSP, to investigate their binding with SPZs, and to compare their ability to take up bystander surface antigens.

Finally, our work provides insights into the development of *Plasmodium* SPZ vaccines. It is known that antibody responses to PfCSP and levels of protection are lower in vaccinated malaria-exposed individuals compared to malaria-naïve individuals ^16^. Therefore, it could be considered to generate transgenic parasites expressing blood stage CD4^+^ T cell epitopes on the surface of SPZs. As such, prior malaria infections should be able to impose a hapten carrier like effect, resulting in a more pronounced antibody response toward SPZ surface antigens like PfCSP and offer better protection in malaria endemic regions.

## Methods

### Mice

C57BL/6 mice, CD28KO^-/-^, MD4, mCherry-rev-mPfCSP-fl, Foxn1 Cre, PfCSP^full-tol^, CST-II, OT-II, PbT-II and Igh^g2A10^ were bred in-house at the Australian National University. All mice were on a C57BL/6 background. Mice used for the experiment were 6 to 10 weeks, and they were age and sex matched for each experiment groups. Mixed female and male mice were used throughout the experiments, and were bred and maintained under specific pathogen free conditions in individually ventilated cages at the Australian National University. Details of all animal strains are summarized in **Supplementary table 1**.

### CST-II TCR transgenic mice and mCherry-rev-mPfCSP-fl mice generation

The method to generate CST-II TCR transgenic mice was reported previously for generating OT-II mice ^43^. The original TCR alpha and beta chain sequences were recovered from scRNA-seq analysis and ligated in to the pES4 and p3A9cbTCR plasmids respectively by Genescript. The resulting plasmids were expanded by *E.coli* culture, extracted by miniprep and digested overnight at 37°C by ClaI-NotI (for pES4) or ApaI-NotI (for p3A9cbTCR) (New England Biolabs). The digested plasmids were run on a 1% TAE agarose gel (with 1:20000 GelRed nucleic acid gel stain, Biotium) and the larger bands were excised for DNA purification. After determining the DNA concentrations by Nanodrop (ThermoFisher Scientific), the DNA sequences of alpha and beta chain were mixed, diluted and microinjected into the mouse embryos. The off-springs were genotyped to check the integration of the TCR transgenes and one founder was produced. This founder can also pass on the transgene to the next generation. The microinjection and mouse transgenic mouse screening was performed by the Phenogenomic Targeting Facility at Australian National University. mCherry-rev-mPfCSP-fl were generated by Ozgene Pty Ltd (Bentley, WA, Australia) via embryonic cell transformation. Briefly the knock-in gene was synthesised and inserted into a plasmid carrying flanking regions for targeting Rosa26 locus. The knock-in gene also carries a neomycin linked with mCherry for selection in embryonic stem cells.

### Bone marrow chimera

WT, PfCSP^T-tol^ and PfCSP^full-tol^ mice were irradiated twice with 500 Rads, and each mouse was intravenously injected with 6-8 million mixed bone marrow cells (50% WT + 50% CST-II) to generate bone marrow chimera mice.

### Plasmodium parasites

*P. berghei* parasites were stored as iRBC stocks in liquid nitrogen. To generate *P. berghei* SPZs, C57BL/6 mice were infected with iRBCs. When parasitemia reached >3% (by Giemsa staining), the mice were anesthetized and placed on an *Anopheles stephensi* mosquito cage to feed for 40 minutes. 18-25 days post blood feed, SPZs were collected by dissecting the mosquito salivary glands, followed by resuspending SPZs in PBS. 2000 SPZs were subsequently injected into mice to establish a blood stage infection and to make new iRBC stocks. These procedures were repeated to maintain all strains of *P. berghei* parasites except for PfCSP-OVA^surface^, which was maintained merely by serial passage through blood stage infection (SPZ of this strain couldn’t establish blood stage infection). Details of all parasite strains are summarized in **Supplementary table 1**.

### Transgenic Plasmodium parasite generation

The method to generate transgenic parasite was reported previously ^44^. The designed DNA sequences were ligated into the Pb268 plasmid which contains the homologous arms of the P. *berghei* chromosome 12, CSP promoter (followed by designed sequences) and DHFR gene (pyrimethamine resistance) under eEF1 promoter ^45^. The resulting plasmids were expanded in E. coli culture, miniprepped and digested overnight at 37°C by PvuI (New England Biolabs) twice. The digested DNA was cleaned up by a miniprep column for future transfection use. To obtain *P. berghei* parasites for transfection, the mice were infected with iRBC and when parasitemia reaches >5 %, the mice were anesthetized by ketamine/xylazine to collect the whole blood in heparin tubes through cardiac puncture. Per 1 mL blood was then diluted into 200 mL RPMI-1640 media with 20% FBS and cultured at 37°C for 24h. Blood smear was performed to confirm the transition of ring stage parasites to schizonts. Then the schizonts were enriched by running the cultured iRBC through a LS column with a 26.5-gauge needle to slow down the flow speed ^46^. After counting, 10^7^ schizonts were resuspended 100ul in human T cell transfection buffer containing 5μg digested DNA for transfection by U-33 program (Lonza).

The transfected iRBCs at schizont stage of the life cycle were immediately intravenously injected into C57BL/6 mice fed by pyrimethamine water. On day7-10 the parasitemia was analyzed, and if it reaches >1%, the iRBC was then cloned by diluting into 0.8 parasite per 100 μL DPBS per mouse and injected into 10-20 mice. The clones that expanded after 7-10 days were further validated by PCR (gDNA was extracted from iRBC) to check for the integration into the chromosome 12 and the presence of the transgenes.

### Immunization

For most experiments, recipient mice were intravenously transferred with 10^4^ antigen-specific Igh^g2A10^ B cells, and with 10^5^ CST-II, OT-II or PbT-II CD4^+^ T cells followed by immunizations. For iRBC immunization, C57BL/6 mice were infected with iRBC stocks. When parasitemia reaches >10% by Giemsa staining, whole blood from these mice was collected into a heparinised tube. 200 μL irradiated (15kRad) iRBC (whole blood) were then intravenously injected into each mouse. For SPZ immunization, each mouse was intravenously injected with 3-6 ×10^4^ irradiated (15kRad) SPZ resuspended in PBS. For rPfCSP-alum immunization, each mouse was intraperitoneally injected with 30μg recombinant PfCSP prepared in Imject™ Alum (7761, ThermoFisher Scientific). In some experiments, mice were intraperitoneally injected with 150μg GK1.5 (BE0003-1, Bio X Cell) to deplete CD4^+^ T cells.

### Flow cytometry

Flow cytometry was performed on a FACS analyser (Fortessa X-20, BD). For spleens, total splenocytes from each mouse were mixed with 5 μL CountBright™ Absolute Counting Beads (C36950, ThermoFisher Scientific) to recover total cell count. ACK lysis buffer (A1049201, ThermoFisher Scientific) treated splenic cells were Fc blocked on ice for 10 minutes and top up by the same volume of 2× concentrated antibody cocktail for another 1 hour on ice, followed by one wash and flow cytometry analysis. For intranuclear staining, surface staining was performed followed by fix/perm by eBioscience™ Foxp3 kit (00-5523-00, ThermoFisher Scientific) and stained for nuclear proteins under room temperature for 1 hour. For TEC analysis, the TEC were prepared as described ^47^. In brief, thymus was cut into fine pieces and digested by liberase for 20 minutes twice (5401119001, Sigma-Aldrich). After wash, surface staining was performed followed by permeabilised using Cytofix/Cytoperm (AB_2869008, BD) and stained for intracellular antibody on ice for 1 hour. For checking the CST-II deletion in thymus, thymus was smashed against a cell strainer and followed by standard surface staining. For SPZ, dissected SPZs were incubated in antibody cocktail on ice for 1 hour followed by flow cytometry analysis. The following reagents are used in staining. CD45.1 (A20, Biolegend); CD45.2 (104, Biolegend); GL7 (GL7, Biolegend); CD38 (90, ThermoFisher Scientific); CD19 (6D5, Biolegend); IgD (11-26c.2a, Biolegend); IgM (II/41, ThermoFisher Scientific); B220 (RA3-6B2, BD); CD4 (GK1.5, Biolegend); Vα2 (B20.1, BD); Vβ3 (KJ25, BD); Igβ (HM79-12, Biolegend); Igκ (187.1, BD); CD44 (IM7, Biolegend); CD62L (MEL-14, BD); CD25 (3C7, Biolegend); FOXP3 (MF-14, Biolegend); CXCR5 (L138D7, Biolegend); PD-1 (RMP1-30, Biolegend); BCL6 (K112-91, BD); NANP9 tetramer (in-house made); Anti-PfCSP (Mab10, in-house made); 7AAD (420404, Biolegend); Zombie Aqua (423102, Biolegend).

### Image flow cytometry and SEM

For image flow cytometry, 10^6^ MACS-enriched B cells or BMDMs were cocultured with 10^4^ sorted GFP-expressing PfCSP-SPZ in a 96-well U bottom plate in complete RPMI-1640 media for at 37 °C for 1 hour, stained for B220, Igβ and PfCSP on ice and analysed by Amnis ImageStream®X MKII (Cytek). Events were collected at 60x objective. GFP^+^ cells were gated for down-stream analysis. The built-in colocalization wizard in IDEAS software was used to analyse the bright detail similarity between the chosen channels. For SEM, B cells and SPZs were cultured as mentioned above, washed in PBS once and fixed at 1% glutaraldehyde for 2 hour at room temperature. After washing with PBS, the fixed cells were transferred onto a Poly-L-Lysine coated coverslip, subjected to a graded ethanol dehydration series, followed by critical point drying and platinum coating. The samples were analysed by ZEISS UltraPlus FESEM at 3kV.

### ELISA

Blood was collected from mice via tail vein and left to clot, and the sera was collected by spinning the blood at 2000 g for 15 minutes. Next, Maxisorp Nunc-Nucleon 96 flat bottom plates (ThermoFisher Scientific) were coated with 5μg/ml recombinant PfCSP overnight at 4 °C, or with 5μg/ml streptavidin overnight at 4 °C followed by 5μg/ml PPPPNPND5 peptide (central repeat of PbCSP) overnight at 4 °C. The following day, plates were washed in wash buffer consisting 0.05% tween20 in PBS, and were blocked for 1 hour before incubating with sera (dilution of 1/100 times, followed by 1 in 3 serial dilutions down the plate) for one hour. After washing, anti-IgG detection antibody conjugated to HRP (5220-0341, Sera Care) was diluted 1/1000 times and was incubated for one hour. The plates were washed, then developed with Peroxidase Substrate Kit (5120-0032, Sera Care) for 10- 15 minutes, and the reaction was stopped by 1% SDS in water. OD405 was then measured. By default, the data was expressed as IC50 from the log(dilution) on the x axis and the OD405 on the y axis, fitting a sigmoidal curve. However, if any sample did not fit in the sigmoidal curve, the area under the curve (AUC) was calculated instead.

### scRNA-seq

Splenocytes were collected on day10 after the immunizations, and splenic CD4^+^ T cells were enriched by the MACS kit (130-104-454, Miltenyi Biotec). The enriched CD4^+^ T cells were stained with antibody cocktails containing flow cytometric and Hashtag antibodies on ice for 30 minutes, washed once and CD4^+^CD44^+^PD-1^+^CXCR5^+^ Tfh cells were sorted. The following Hashtag antibodies were used. TotalSeq-C anti-mouse Hashtag 1, 2, 3, 4 and 5 (M1/42 and 30-F11, Biolegend). 10^4^ pooled Tfh cells from different mice were loaded onto each lane for of the 10X Chromium platform (10X Genomics). Library preparation was completed by Biomedical Research Facility at the Australian National University following the recommended protocols. Libraries were sequenced using the NovaSeq6000 (Illumina). The 10× Cell Ranger package was used to process transcript, CITE-seq and VDJ libraries for downstream analysis.

### Bioinformatic analysis for scRNA-seq

The analysis of scRNA-seq datasets has been described in our previous publication ^48^. In brief, transcripts and CITE-seq library outputs were loaded into the Seurat package ^49^ for unwanted cell removal, clustering, annotation and visualization such as UMAP, violin plot, feature plot and dot plot. VDJ analysis was performed using the functions in the scRepertoire and ClustIRR R packages ^50^.

### Statistical analysis

Statistical analysis was performed by Prism software (GraphPad). For plotting cell numbers on a log scale, the cell counts of all samples were added by 1 if any value was originally 0 and log10 transformed. All statistics were showed as mean ± standard error of the mean. All data were assumed Gaussian distributed thus comparisons between two groups were performed by two- tailed parametric t-test and multiple comparisons were performed by One-Way or Two-Way ANOVA post hoc Dunnett, Sidak or Tukey test using the default settings of Prism. P values ≤ 0.05 were considered significant and specified in the paper.

## Data availability

The original and processed single-cell RNA-seq data have been deposited at Gene Expression Omnibus (GSE274341).

## Code availability

The code for bioinformatic analysis is available at https://doi.org/10.6084/m9.figshare.26526313.

## Supporting information

Gao et al. Supplementary Information

## Acknowledgements

We thank support from H. Vohra and M. Devoy of the imaging and cytometry facility at the Australian National University. We acknowledge the instruments and expertise of Microscopy Australia at the Centre for Advanced Microscopy, the Australian National University, a facility enabled by NCRIS and university support. We thank Siming Man lab for providing BMDMs. X.G. is supported by a PhD studentship from the Australian National University. Work in the Cockburn lab is supported by an NHMRC Investigator Grant to I.A.C. (GNT2008648). The funders had no role in the drafting of the manuscript or the decision to publish.

## Contribution

Study design: I.A.C., X.G., H.A.M..

Transgenic mice: X.G., I.A.C., D.F., D.S..

Transgenic parasites: X.G., K.W., F.F., A.J.S..

Mouse experiments: X.G., H.A.M., K.W., H.G.K., A.F. L., P.C., L.X., L.B., D.C.T., D.H.D.G..

Imaging experiments: X.G., J.L., M.R., K.P.. Bioinformatics: X.G., A.D..

Manuscript writing: I.A.C., X.G..

## Corresponding author

Correspondence to Ian A. Cockburn

## Ethics declarations

All animal procedures were approved by the Animal Experimentation Ethics Committee of the Australian National University (Protocol number: 2019/36 and 2022/36). All research involving animals was conducted in accordance with the National Health and Medical Research Council’s

Australian Code for the Care and Use of Animals for Scientific Purposes and the Australian Capital Territory Animal Welfare Act 1992.

