## Supplementary material for "B cells targeting parasites capture spatially linked antigens to secure T cell help": Gao et al. Supplementary Information

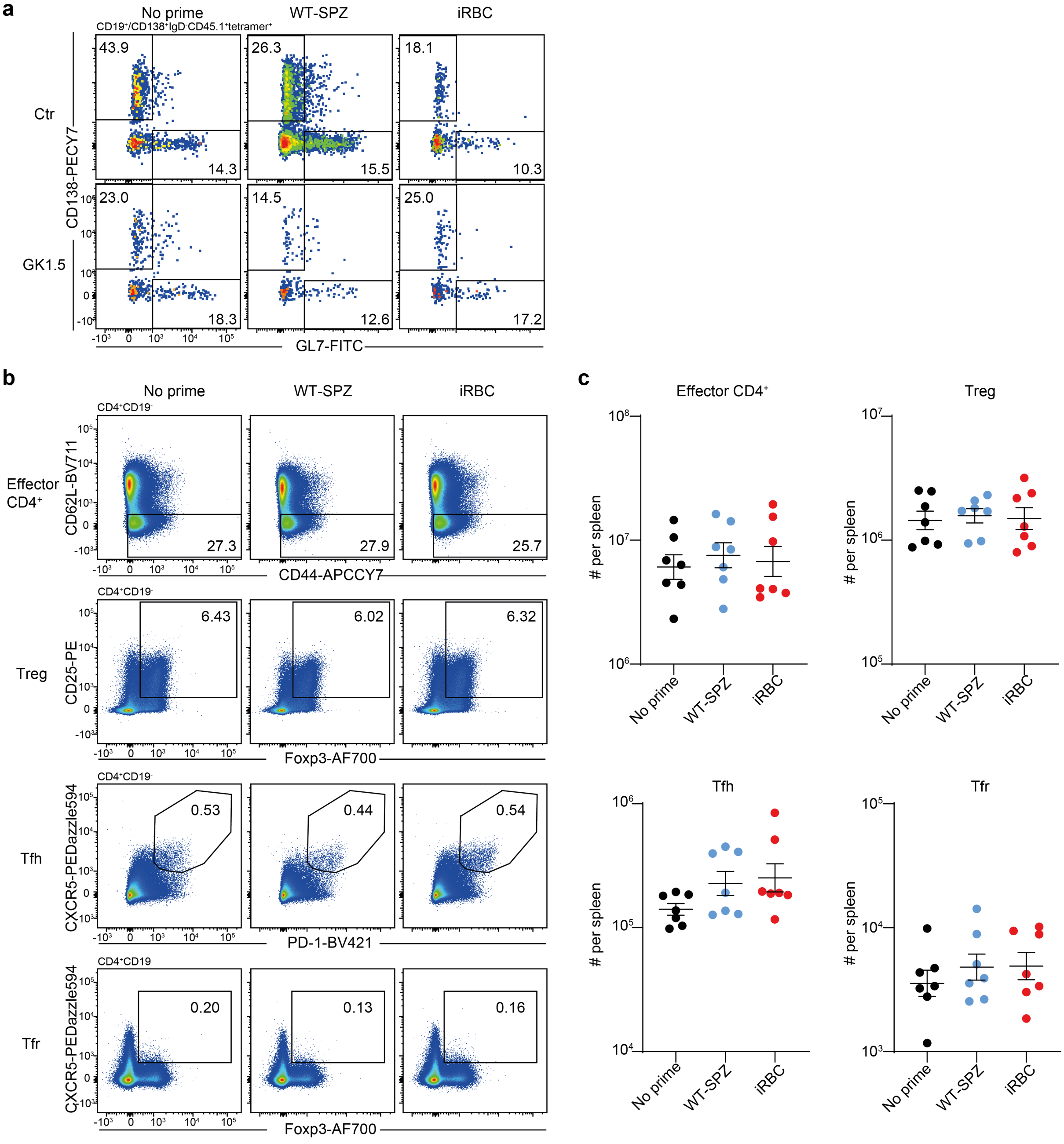


**Extended data figure 1. No detectable differences in the formation of CD4+ T cell subsets between iRBC and WT-SPZ primed mice.**

WT mice were untreated, or primed by irradiated WT-SPZ or iRBC, followed by Igh$^{g2A10}$ B cell transfer w/wo 150µg GK1.5 treatment, and boost by PfCSP-SPZ. Spleens were analysed (n≥3). **a**, Representative FACS plots showing the formation of Igh$^{g2A10}$ PB/PC (CD138$^{+}$) and GCB (GL7$^{+}$) cells. **b**, Representative FACS plots and **c**, statistics showing the formation of indicated CD4$^{+}$ T cell subsets. Regulatory T cells (Treg). Regulatory follicular T cells (Tfr). Results were pooled from two independent experiments for c.

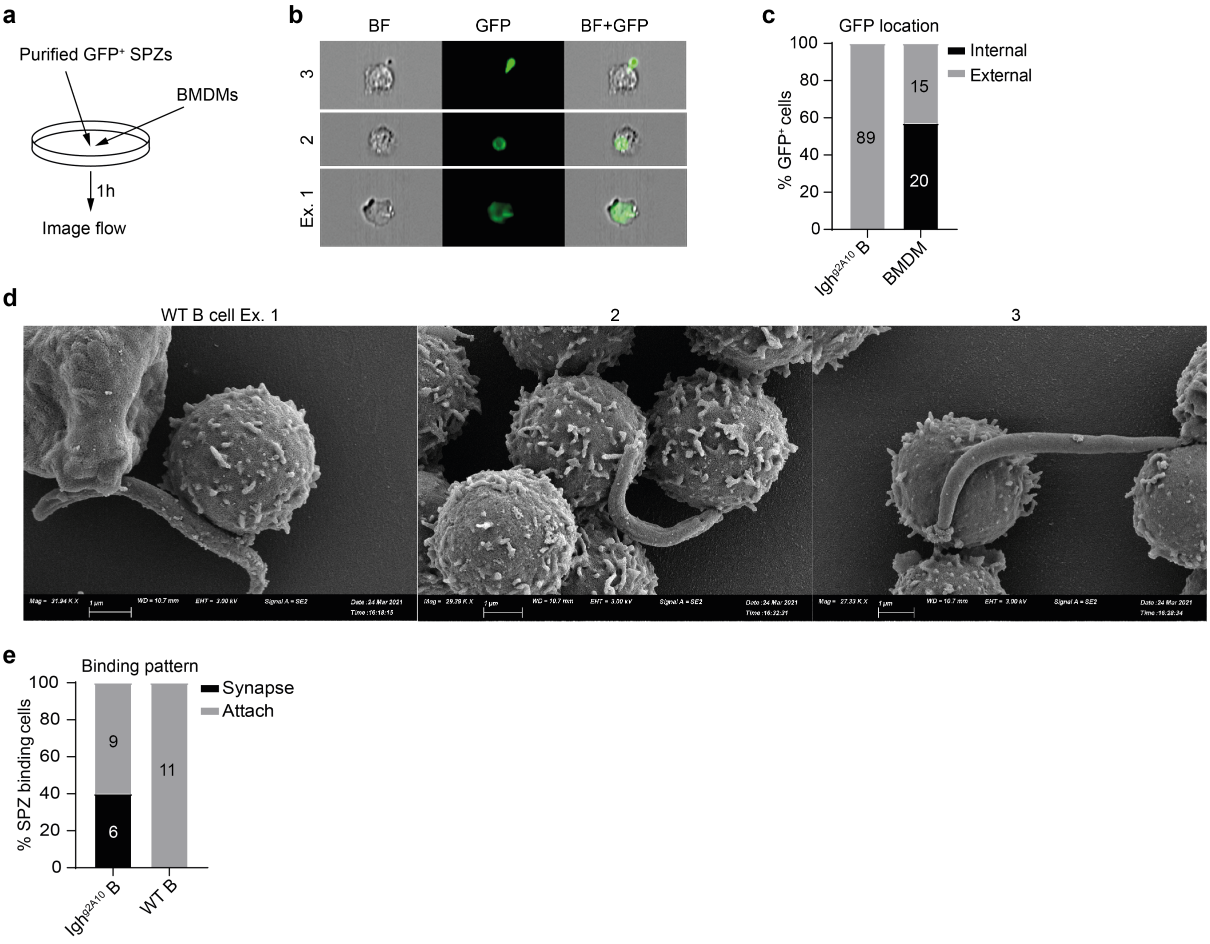


**Extended data figure 2. BMDM can phagocytose SPZs in vitro.**

**a-b**, $10^6$ BMDMs were incubated with $10^4$ GFP$^+$ PfCSP-SPZ per well at 37°C for 1 hour, followed by image flow analysis. **a**, Experiment design and **b**, representative images for GFP$^+$ BMDMs. **c**, Statistics showing the numbers (within bars) and percentages of images with indicated GFP location. **d**, Representative SEM images of SPZ-interacting WT B cells. **e**, Statistics showing the numbers (within bars) and percentages of images with indicated SPZ binding pattern. Results were representative of two independent experiments for b,d, and pooled from two independent experiments for c,e.

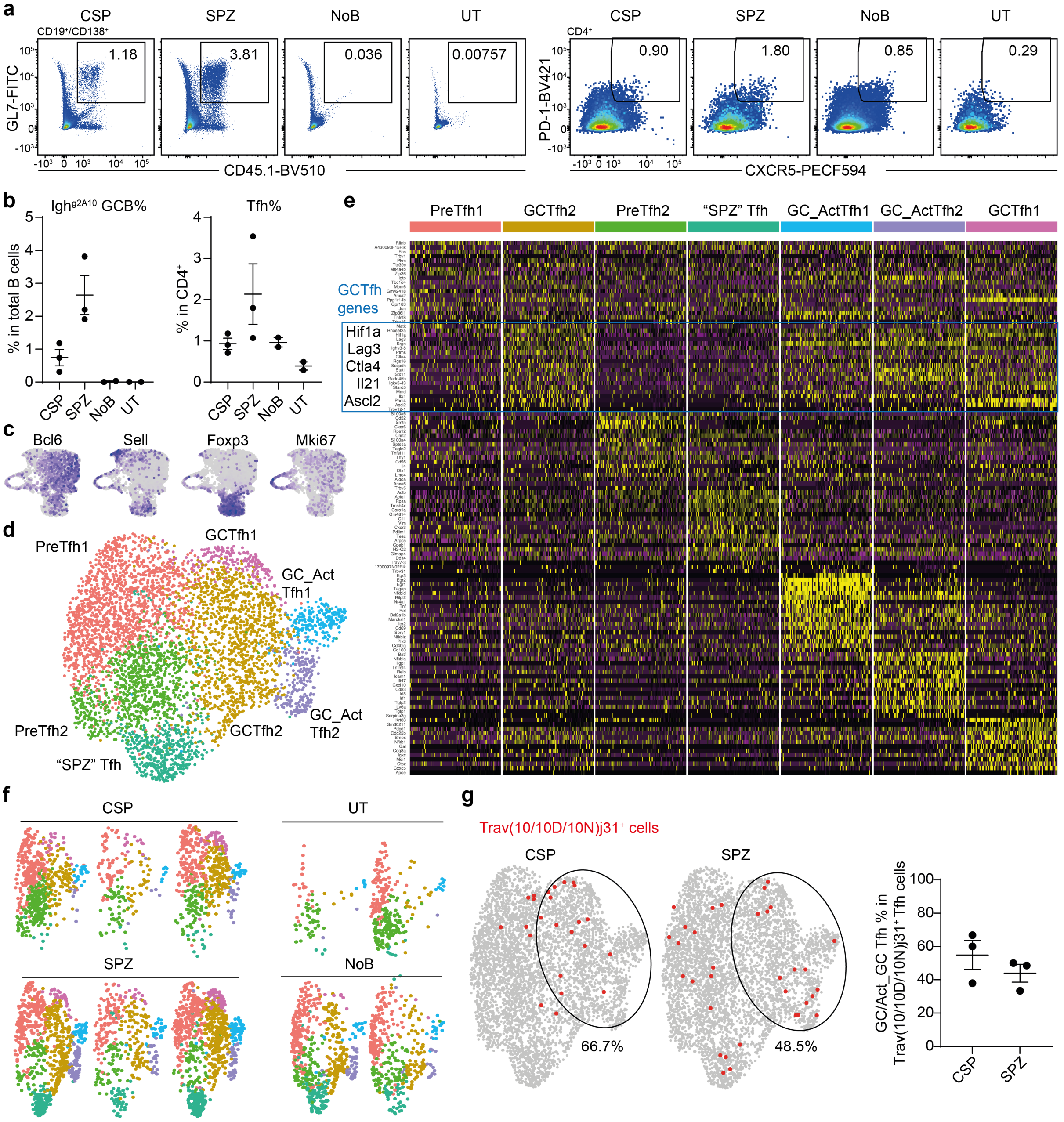


**Extended data figure 3. scRNA-seq analysis for Igh$^{g2A10}$-helping Tfh cells.**

MD4 mice were untreated or transferred with Igh$^{g2A10}$ B cells, followed by rPfCSP-alum, PfCSP-SPZ immunization or left untreated, and spleens were collected. Tfh cells (CD4$^{+}$CD44$^{+}$PD-1$^{+}$CXCR5$^{+}$) were FACS-purified, hashtaged and pooled from scRNA-seq. **a**, Representative FACS plots and **b**, statistics showing the formation of Igh$^{g2A10}$ GCB cells and Tfh cells (same samples for scRNA-seq). **c**, Feature plots for the expression of indicated genes. **d**, UMAP of reclustered Bcl6$^{+}$ Tfh cells. **e**, Heatmap of signature genes expression in Bcl6$^{+}$ Tfh sub-clusters. **f**, UMAPs of reclustered Bcl6$^{+}$ Tfh cells separated by mouse. **g**, Representative UMAPs and statistics showing proportion of Trav(10/10D/10N)j31$^{+}$ cells that belongs to GC Tfh clusters (circled area).

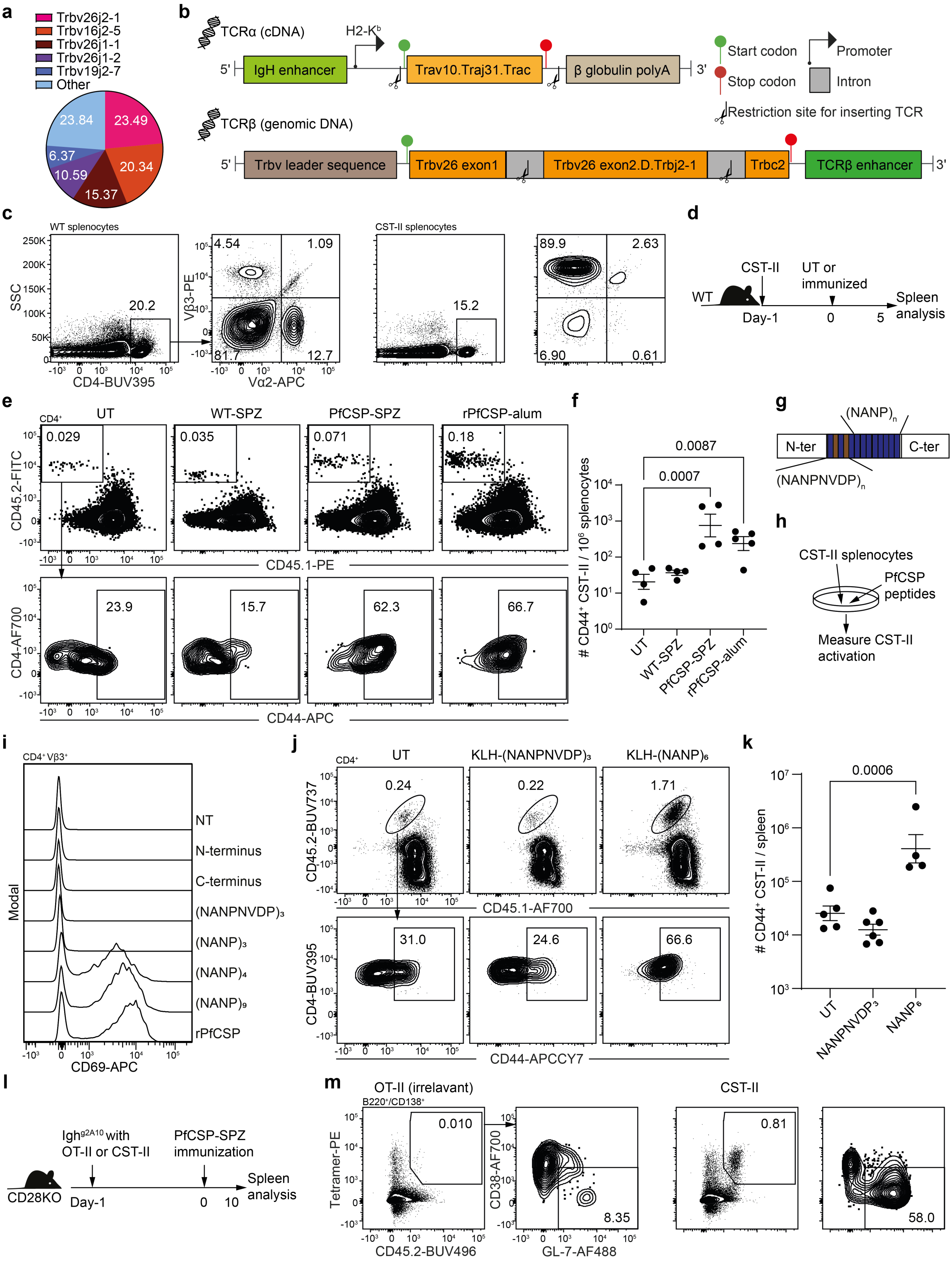


**Extended data figure 4. Generation and characterization of TCR transgenic mice CST-II.**

**a**, Trbv/j proportions of TCRβs paired with Trav10j31. **b**, DNA constructs to generate CST-II mice. **c**, Representative FACS plots showing the expression of Trbv26 (vβ3) and irrelevant vα2 on $CD4^+$ T cells in WT and CST-II mice. d-f, $10^5$ CST-II $CD4^+$ T cells were transferred into WT mice followed by indicated immunizations, spleens were analyzed. **d**, Experiment design. **e**, Representative FACS plots and **f**, statistics showing the formation of effector CST-II cells. **g**, Diagram of PfCSP structure. **h-i**, CST-II splenocytes were cultured with 1μg /ml indicated peptides followed by FACS analysis. **h**, Experiment design and **i**, representative FACS plots showing the expression of CD69. **j-k**, $10^5$ CST-II $CD4^+$ T cells were transferred into WT mice followed by indicated immunizations, spleens were analyzed**.** **j**, Representative FACS plot and **k**, statistics showing the formation of effector CST-II cells. **l-m**, $10^4$ $Igh^{g2A10}$ B cells were co-transferred with $10^5$ OT-II or CST-II $CD4^+$ T cells into $CD28^{-/-}$ mice followed by PfCSP-SPZ immunization, and spleens were analysed. **l**, Experiment design and **m**, representative FACS plots showing the formation of $Igh^{g2A10}$ GCB cells. Results were representative for ≥five independent experiments for c,m, for two independent experiments for i, and pooled from two independent experiments for f,k. P values were calculated by One-Way ANOVA with Dunnett multiple comparisons test for f,k.

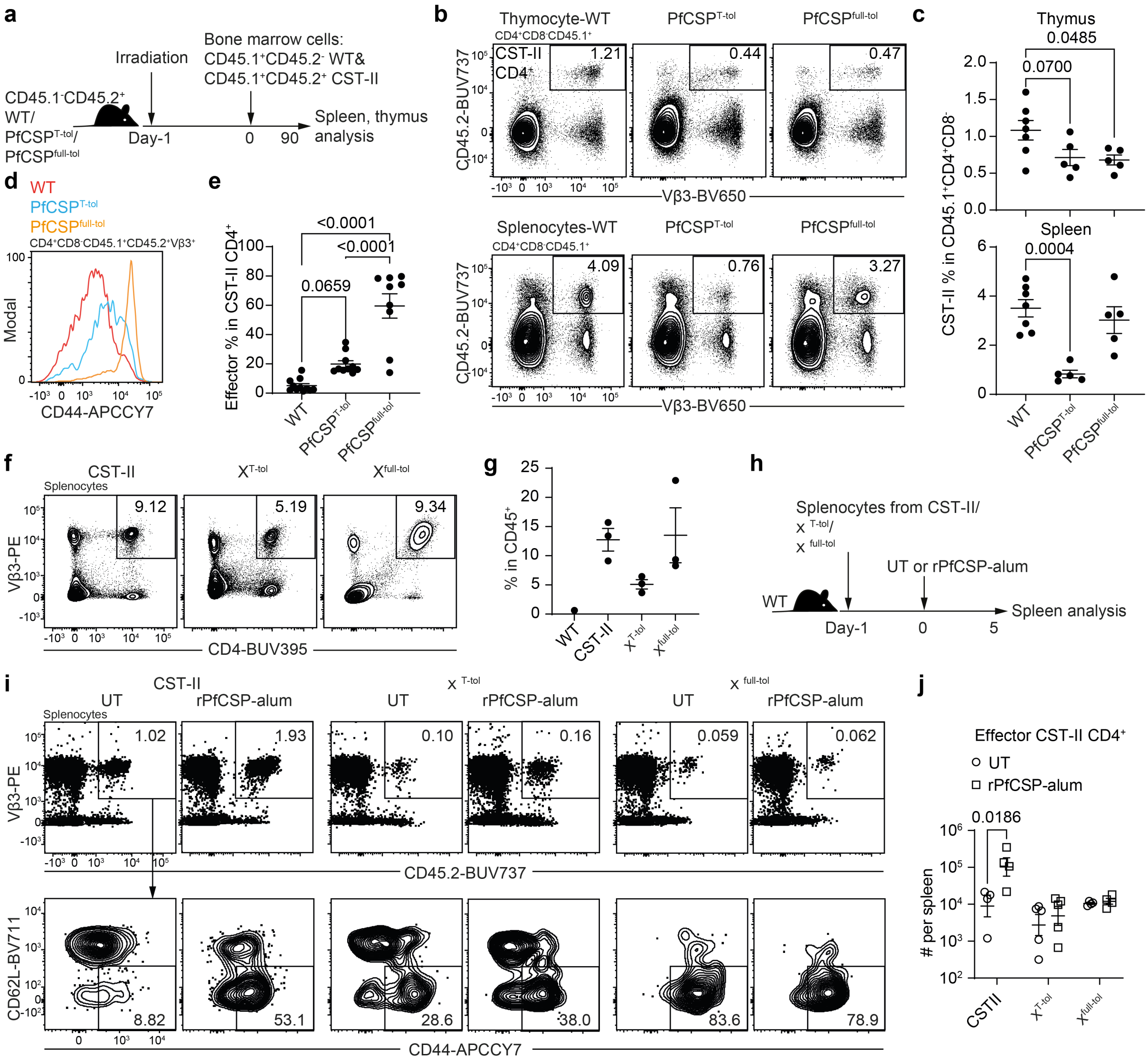


**Extended data figure 5. Assessing $CD4^+$ T cell tolerance in $PfCSP^{T\text{-}tol}$ and $PfCSP^{full\text{-}tol}$ mice.**

**a-e**, WT, $PfCSP^{T\text{-}tol}$ and $PfCSP^{full\text{-}tol}$ mice were sub-lethally irradiated and transferred with 5-10 million mixed bone marrow cells from CST-II (50%) and WT (50%) mice. Thymus and spleens were analysed. **a**, Experiment design. **b**, Representative FACS plots and **c**, statistics comparing the formation of CST-II $CD4^+$ T cells in indicate tissues and mice. **d**, Representative FACS plots showing the CD44 expression on CST-II $CD4^+$ T cells from indicated mice. **e**, Statistics showing the percentages of effector CST-II $CD4^+$ T cells from indicated mice. **f-j**, $PfCSP^{T\text{-}tol}$ and $PfCSP^{full\text{-}tol}$ mice were crossed to CST-II background and the splenocytes from the resulting mice were analysed, or transferred into WT recipients followed by rPfCSP-alum immunizations and spleens were analysed. **f**, Representative FACS plots and **g**, statistics showing the formation of CST-II $CD4^+$ T cells in indicated mice. **h**, Experiment design. **i**, Representative FACS plots and **j**, statistics showing the formation of effector CST-II $CD4^+$ T cells w/wo immunization. Results were representative for two independent experiments for c, pooled from two independent experiments for e,g,j. P values were calculated by One-Way ANOVA with Dunnett multiple comparisons test for c, with Tukey multiple comparisons test for e, and by Two-Way ANOVA with Šidák multiple comparisons for j.

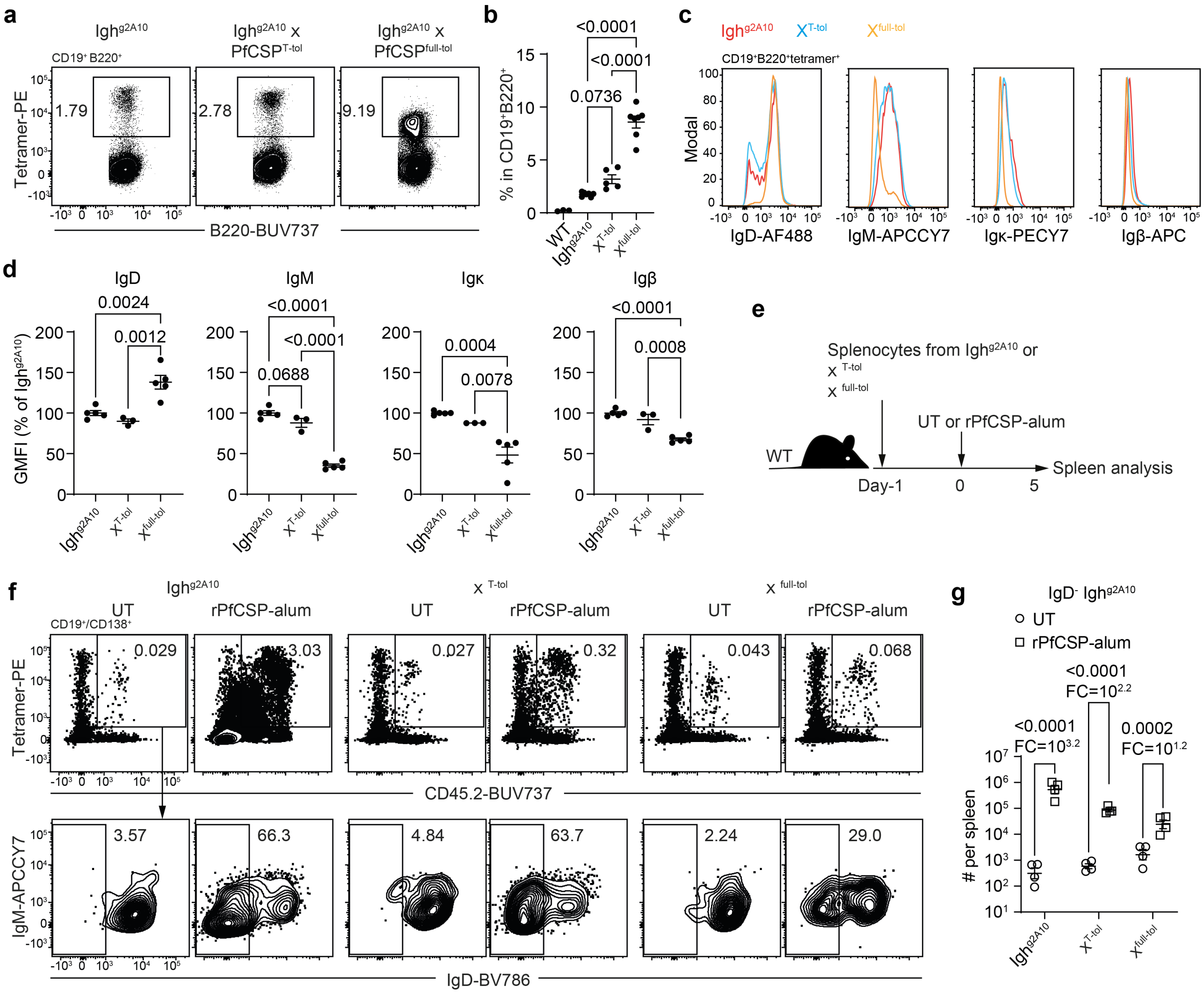


**Extended data figure 6. Assessing B cell tolerance to PfCSP in PfCSP$^{\text{T-tol}}$ and PfCSP$^{\text{full-tol}}$ mice.**

PfCSP$^{\text{T-tol}}$ and PfCSP$^{\text{full-tol}}$ mice were crossed to Igh$^{\text{g2A10}}$ background and the splenocytes from the resulting mice were analysed, or transferred into WT mice followed by rPfCSP-alum immunization, and spleens were analysed. **a**, Representative FACS plots and **b**, statistics showing the proportions of PfCSP tetramer-specific B cells (n≥2). **c**, Representative FACS plots and **d**, statistics showing the expression of indicated markers on tetramer$^{+}$ B cells (n≥1). **e**, Experiment design. **f**, Representative FACS plots and **g**, statistics showing the formation of IgD$^{-}$ Igh$^{\text{g2A10}}$ B cells w/wo immunization (n=2). Results were pooled from two independent experiments for b,d,g. P values were calculated by One-Way ANOVA with Tukey multiple comparisons test for b,d, and by by Two-Way ANOVA with Šidák multiple comparisons for g.

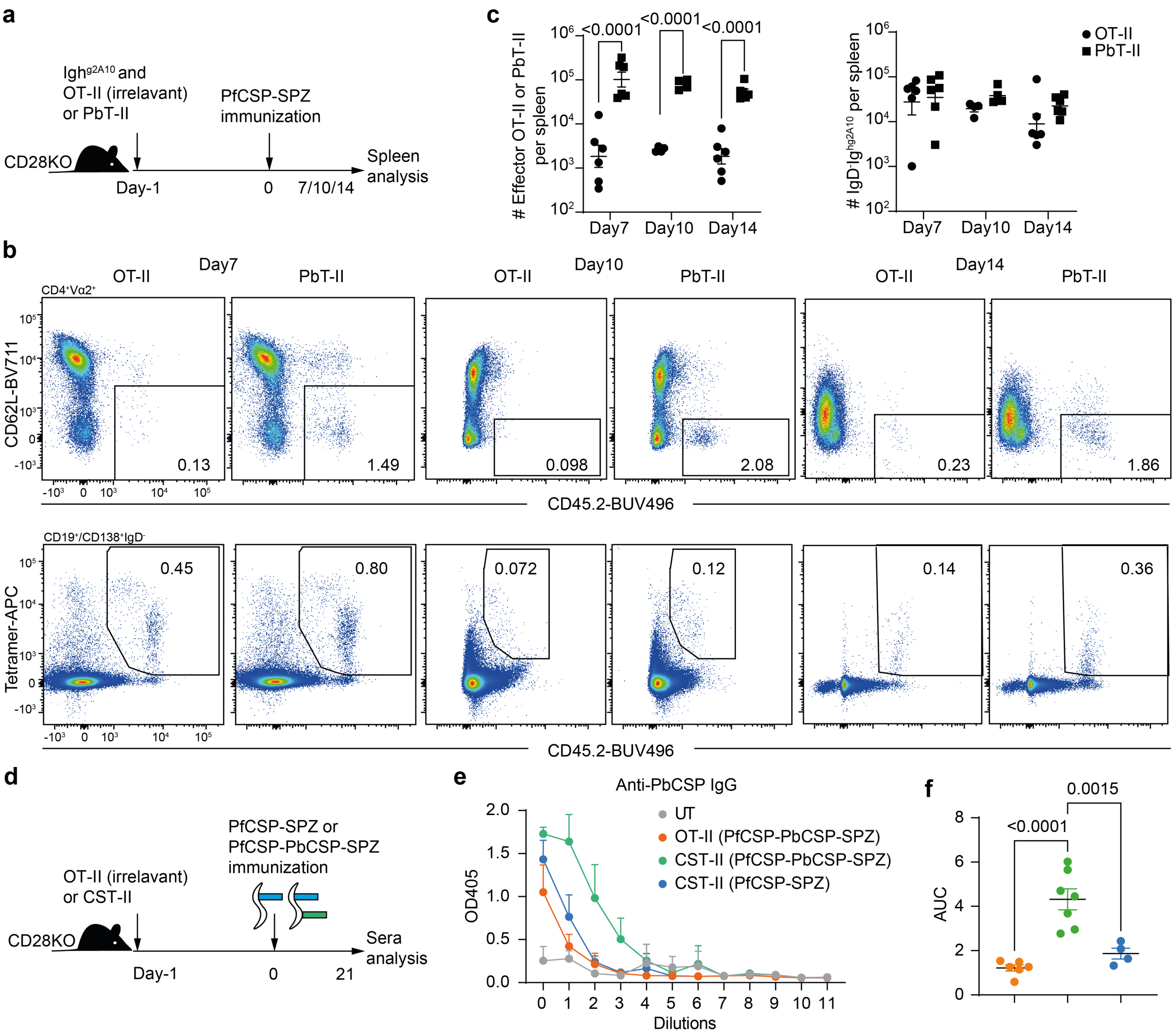


**Extended data figure 7. Inter-molecular help for CSP-specific B cells comes from CD4$^+$ T cells specific for antigens on the surface but not interior of SPZ.**

**a-c**, CD28$^{-/-}$ mice were transferred with Igh$^{g2A10}$ B cells with OT-II or PbT-II cells, followed by PfCSP-SPZ immunization, and spleens were analysed (n≥3). **a**, Experiment design. **b**, Representative FACS plots and **c**, statistics showing the formation of effector OT-II or PbT-II CD4$^+$ T cells and IgD$^-$ Igh$^{g2A10}$ B cells. **d-f**. CD28$^{-/-}$ mice were transferred with OT-II or CST-II CD4$^+$ T cells, followed by PfCSP-SPZ or PfCSP-PbCSP-SPZ immunization. The mice were bleed to measure anti-PbCSP IgG (n≥2). **d**, Experiment design. **e**, OD405 for serial diluted sera and **f,** statistics showing the titers of anti-PbCSP IgG. Results were pooled from two independent experiments for c,f. P values were calculated by Two-Way ANOVA with Šidák multiple comparisons for c, and by One-Way ANOVA with Tukey multiple comparisons test for f.

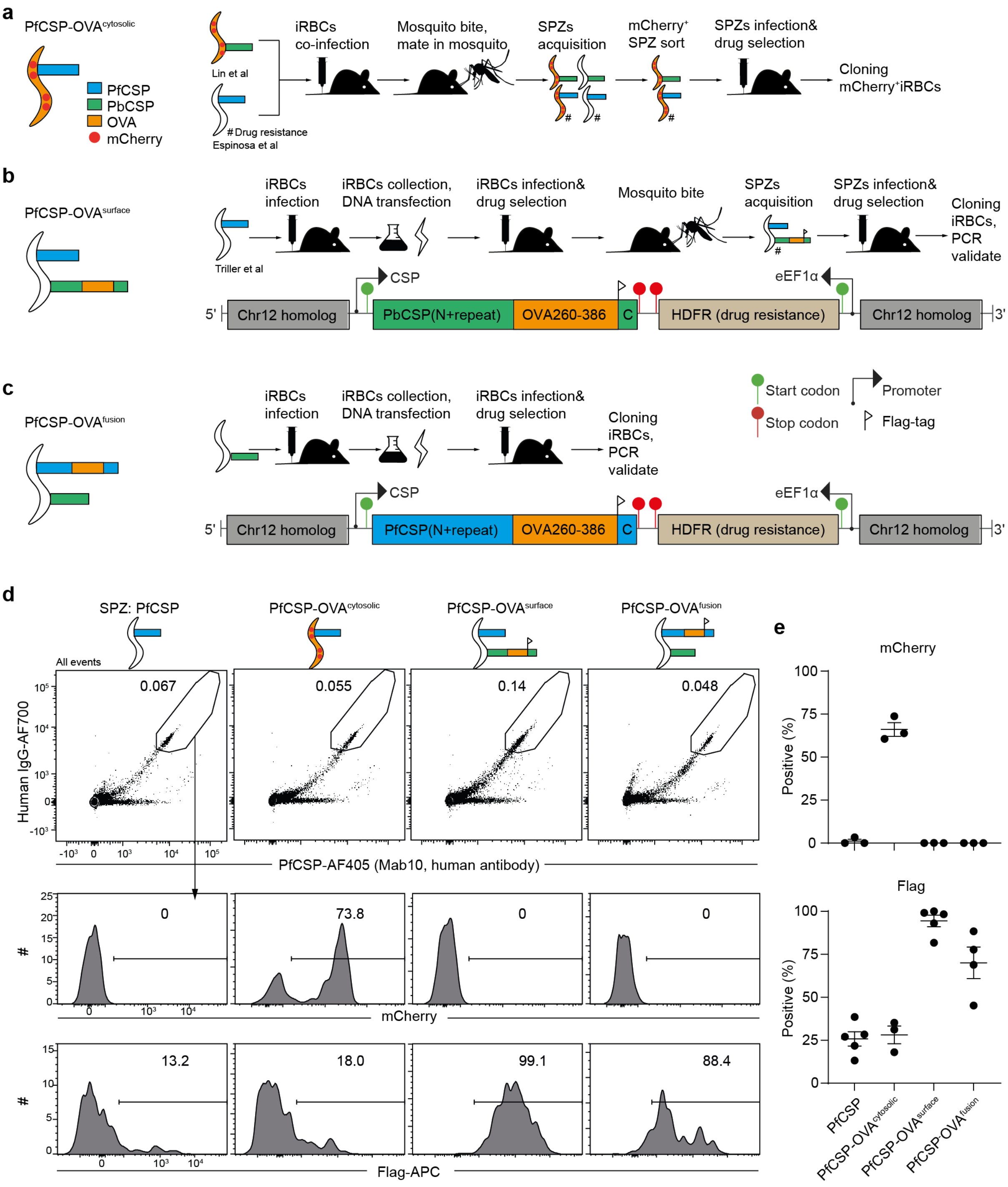


**Extended data figure 8. Generation and characterization of transgenic *P.berghei* SPZ expressing OT-II epitopes on different locations.**

Diagram showing the DNA constructs and generation route of SPZ expressing full length OVA or $OVA_{260\text{-}386}$ **a**, in cytosol (PfCSP-OVA$^{cytosolic}$), **b**, on the surface by fused within PbCSP molecule (PfCSP-OVA$^{surface}$), or **c**, fused within PfCSP molecule (PfCSP-OVA$^{fusion}$). d-e, Transgenic sporozoites were collected from the mosquito salivary glands and phenotyped by FACS (n=1, pooled from ~150 mosquitos). **d**, Representative FACS plots and **e**, statistics showing the expression of indicated markers. Results were pooled from three independent experiments for e.

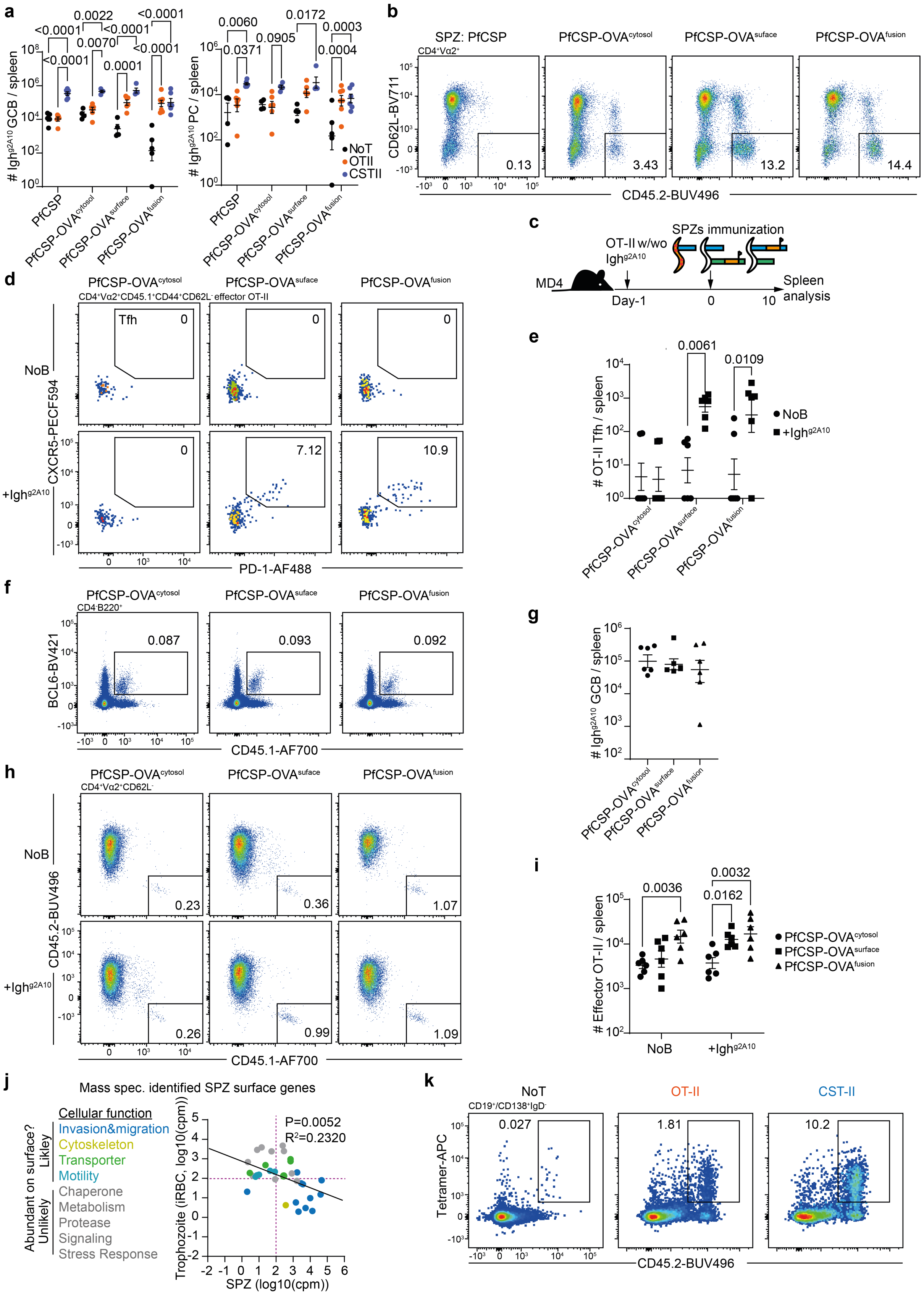


**Extended data figure 9. Ighg2A10 B cells can present surface but not interior antigens during SPZ immunization.**

**a-b**, $CD28^{-/-}$ mice were transferred with $Igh^{g2A10}$ B cells w/wo OT-II or CST-II $CD4^+$ T cells, immunized with transgenic SPZs, and spleens were analysed. **a**, Statistics showing the formation of $Igh^{g2A10}$ GCB and PC. **b**, Representative FACS plots showing the formation of effector OT-II cells upon indicated SPZ immunization. **c-i**, MD4 mice were transferred with $10^5$ OT-II $CD4^+$ T cells w/wo $10^4$ $Igh^{g2A10}$ B cells, and immunized by transgenic SPZs. Spleens were collected for FACS analysis (n=3). **c**, Experiment design. **d**, Representative FACS plots and **e**, statistics showing the formation of OT-II Tfh cells. **f**, Representative FACS plots and **g**, statistics showing the formation of $Igh^{g2A10}$ GCB cells. **h**, Representative FACS plots and **i,** statistics showing the formation of effector OT-II $CD4^+$ T cells. **j**, mRNA expression of mass spectrum identified SPZ surface proteins and their cellular function as an indication for surface abundance. We identified 32 *P. berghei* genes that were homologous to *P. falciparum* genes, and had annotated cellular function. **k**, $CD28^{-/-}$ mice were transferred w/wo OT-II or CST-II $CD4^+$ T cells, immunized by PfCSP-$OVA^{suface}$-SPZ, transferred with $Igh^{g2A10}$ B cells 2 day after immunization, and spleens were analysed on day10 post immunization (n≥2). Representative FACS plots showing the formation of $IgD^-$ $Igh^{g2A10}$ B cells. Results were pooled from three independent experiments for b and two independent experiments for e,g,i. P values were calculated by Two-Way ANOVA with Tukey multiple comparisons test for a,i, and with Šidák multiple comparisons for e.

**Supplementary table1. Mouse and parasite strains**

| Name | Phenotype | Source |
|---|---|---|
| *Mice* | | |
| C57BL/6 (B6) | An inbred wildtype mouse strain. Carries the CD45.2 congenital pan-leukocyte marker. | Bred in House, from the Jackson Laboratories (USA). |
| B6N-CD45.1 | An inbred wildtype mouse strain. Carries the CD45.1 congenital pan-leukocyte marker. | Bred in House, from the Jackson Laboratories (USA). |
| $CD28^{-/-}$ | CD28 is required for T cell activation. This strain has defective $CD4^+$ and $CD8^+$ T cell responses. | Gift from Carola Vinuesa Lab. Reference [51]. |
| MD4 | BCR transgenic mice in which all the B cells are specific for hen egg lysozyme. | Gift from Carola Vinuesa Lab. Reference [52]. |
| $Igh^{g2A10}$ | BCR knock-in mice in which all B cells express the heavy chain of 2A10 antibody that binds to PfCSP $NANP_{repeat}$. | Generated by Ozgene Pty Ltd (WA, Australia). Reference [18]. |
| OT-II | TCR transgenic mice in which $CD4^+$ T cells are specific for the $Ovalbumin_{323-339}$. | Reference [53]. |
| PbT-II | TCR transgenic mice in which $CD4^+$ T cells are specific for the *P. berghei* Hsp90 antigen. | Gift from William Heath Lab. Reference [54]. |
| CST-II | TCR transgenic mice in which $CD4^+$ T cells are specific for the PfCSP $NANP_{repeat}$. | Generated in this study. Generated by Phenogenomic Targeting Facility, Australian National University. |
| mCherry-rev-mPfCSP-fl | Ubiquitous mCherry expression. Cre induces mCherry loss and mPfCSP expression. | Generated in this study. Generated by Ozgene. |
| Foxn1 Cre | Cre controlled by Foxn1 promoter, expressed in thymus epithelia cells. | Gift from Daniel Gray Lab. Reference [55]. |
| $PfCSP^{full-tol}$ | Ubiquitous mPfCSP expression. | Generated in this study. Generated by Ozgene by crossing mCherry-rev-mPfCSP-fl with PGK Cre mice. |
| *P. berghei parasites* | | |
| WT-SPZ/iRBC | Wild type *P. berghei ANKA* strain | Gift from Fidel Zavala Lab. |
| PfCSP-SPZ | Replacement of PbCSP with PfCSP. Has DHFR drug selection marker. | Reference [19] |
| $GFP^+$ PfCSP-SPZ | Replacement of PbCSP with PfCSP. Express GFP. | Reference [18] |
| PfCSP-SPZ | Replacement of PbCSP with PfCSP. No drug selection marker. | Reference [56] |
| PbCSP-PfCSP-SPZ | Co-expressing PbCSP and PfCSP on the surface of SPZ. | Reference [28] |
| PfCSP-$OVA^{cytosol}$-SPZ | Replacement of PbCSP with PfCSP and express OVA-mCherry fusion protein cytosolically under same construct. | Generated in this study by crossing the OVA-mCherry expressing *P.berghei* [57] with PfCSP expressing *P.berghei* [19]. |
| PfCSP-$OVA^{surface}$-SPZ | Express PfCSP and PbCSP-$OVA_{260-385}$-Flag fusion protein which is under the CSP promoter inserted at chr12 non-coding locus. | Generated in this study by transfecting the *P. berghei* that has PbCSP replaced by PfCSP [56]. |
| PfCSP-$OVA^{fusion}$-SPZ | Express PbCSP and PfCSP-$OVA_{260-385}$-Flag fusion protein which is under the CSP promoter inserted at chr12 non-coding locus. | Generated in this study by transfecting wild type *P. berghei*. |
